# Single-nucleus transcriptomics reveals functional compartmentalization in syncytial skeletal muscle cells

**DOI:** 10.1101/2020.04.14.041665

**Authors:** Minchul Kim, Vedran Franke, Bettina Brandt, Elijah D. Lowenstein, Verena Schöwel, Simone Spuler, Altuna Akalin, Carmen Birchmeier

**Affiliations:** Developmental Biology/Signal Transduction, Max Delbrueck Center for Molecular Medicine, Berlin, Germany; Bioinformatics, Berlin Institute for Medical Science Biology, Max Delbrueck Center for Molecular Medicine, Berlin, Germany; Muscle Research Unit, Experimental and Clinical Research Center, Charité Medical Faculty and Max Delbrueck Center for Molecular Medicine, Berlin, Germany

## Abstract

Syncytial skeletal muscle cells contain hundreds of nuclei in a shared cytoplasm. We investigated nuclear heterogeneity and transcriptional dynamics in the uninjured and regenerating muscle using single-nucleus RNA-sequencing (snRNAseq) of isolated nuclei from muscle fibers. This revealed distinct nuclear subtypes unrelated to fiber type diversity, completely novel subtypes as well as the expected ones at the neuromuscular and myotendinous junctions. In fibers of the *Mdx* dystrophy mouse model, new subtypes emerged, among them nuclei expressing a repair signature that were also abundant in the muscle of dystrophy patients, and a nuclear population associated with necrotic fibers. Finally, modifications of our approach revealed the compartmentalization in the rare and specialized muscle spindle. Our data identifies new nuclear compartments of the myofiber and defines a molecular roadmap for their functional analyses; the data can be freely explored on the MyoExplorer server (https://shiny.mdc-berlin.net/MyoExplorer/).

## Introduction

All cells need to organize their intracellular space to properly function. In doing so, cells employ various strategies like phase separation, polarized trafficking, and compartmentalization of metabolites^1-4^. Syncytial cells face an additional challenge to this fundamental problem because individual nuclei in the syncytium can potentially have distinct functions and express different sets of genes. An outstanding example of this is the skeletal muscle fiber, a syncytium containing hundreds of nuclei in very large cytoplasm that possesses functionally distinct compartments. The best documented compartment is located below the neuromuscular junction (NMJ), the synapse formed between motor neurons and muscle fibers. NMJ form in a narrow central region of the fiber, and are characterized by the enrichment of proteins that function in the transmission of the signal provided by motor neurons to elicit muscle contraction^5-9^. Motor neurons are known to instruct myonuclei at the synapse to express genes that function in synaptic transmission. Another specialized compartment is located at the end of the myofibers where they attach to the tendon, allowing force transmission. Many cell adhesion and cytoskeletal proteins are known to be enriched at the MTJ^10,11^. However, little is known about the transcriptional characteristics of MTJ myonuclei and to date only a few genes like *Col22a1, Ankrd1* and *LoxL3* were reported to be specifically expressed at the mammalian MTJ^12-14^. Previous studies have reported that the diffusion of transcripts and proteins in myofibers is limited, and indeed specific transcripts and proteins associated with the NMJ and MTJ appear to diffuse little inside the fiber^5,11,15,16^. Therefore, locally regulated transcription plays an important role in establishing functional compartments in the muscle. In addition, stochastic transcription of particular genes has been reported in myofibers, but it is unknown whether this reflects differences in myonuclear identities^17^. Since a systematic analysis is currently lacking, we neither know the extent of myonuclear heterogeneity nor can we assess whether additional myonuclear types exist beyond those at the NMJ and MTJ. Such knowledge may provide insight into how skeletal muscle cells orchestrate their many functions.

Previous studies on gene expression in the muscle relied on the analysis of selected candidates by *in situ* hybridization or on profiling the entire muscle tissue. The former is difficult to scale up, whereas the latter averages the transcriptomes of all nuclei. More recently, several studies have used single-cell approaches to reveal the cellular composition of the entire muscle tissue^18-20^. However, these approaches did not sample the syncytial myofibers. Single-nucleus RNA-Seq (snRNAseq) using cultured human myotubes failed to detect transcriptional heterogeneity among nuclei^21^, underscoring the importance of studying the heterogeneity in an *in vivo* context where myofibers interact with surrounding cell types.

## Results

### Single-nucleus RNA-Seq analysis of uninjured and regenerating muscles

We genetically labeled mouse myonuclei by crossing a myofiber-specific Cre driver (*HSA-Cre*) with a Cre-dependent H2B-GFP reporter. H2B-GFP is deposited at the chromatin, which allows us to isolate single myonuclei using flow cytometry. Nuclei of regenerating fibers were also efficiently labeled 7 days after cardiotoxin-induced injury (7 days post injury; 7 d.p.i.) (Supplementary Fig. 1a; note that nuclei in uninjured and regenerating fiber locate peripherally and centrally, respectively^22^). We confirmed the efficiency and specificity of the H2B-GFP labeling (Supplementary Fig. 1a-1c). H2B-GFP was absent in endothelia (Cd31+), Schwann cells (Egr2+), tissue resident macrophages (F4/80+) and muscle stem cells (Pax7+) (Supplementary Fig. 1c); nuclei of these diverse cell types lie outside the fiber and together make up around 50% of all nuclei in the tissue.

We next established a protocol for the rapid isolation of myonuclei. Conventional methods involve enzymatic dissociation of muscle fibers at 37°C, which can cause secondary changes in gene expression^23-25^. We used a procedure that took 20 minutes from dissection to flow cytometry, employing fast mechanical disruption on ice. Indeed, our subsequent analysis indicated that this protocol avoided the expression of stress-induced genes (see Methods).

For snRNAseq profiling, we used the CEL-Seq2 technology^26^, a low throughput plate-based method with high gene detection sensitivity^27^. Considering only exonic reads and genes detected in at least 5 nuclei, we detected 1000-2000 genes per nucleus (Supplementary Fig. 2a-2b). Median mitochondrial read thresholds were 1.3% or less in all samples used in this study (Supplementary Fig. 2c). We analyzed nuclei from uninjured (1,591 nuclei) and regenerating *tibialis anterior* (TA) muscle (7 and 14 d.p.i., 946 and 1,661 nuclei, respectively). Uniform Manifold Approximation and Projection (UMAP) analysis of these datasets revealed heterogeneity among myonuclei (Fig. 1a). All nuclei expressed high levels of *Ttn*, a pan-muscle marker (Fig. 1b). The TA muscle contains three different fiber types (IIA - intermediate, IIB - very fast and IIX - fast) that express distinct myosin genes. The largest cluster, bulk myonuclei, could be sub-divided into nuclei from distinct fiber types (Fig. 1b); *Myh1*-(IIX; lower left part of the cluster) or *Myh4* (IIB; upper right part of the cluster)-positive nuclei were most abundant and present roughly in a ratio of 1:1. *Myh2* (IIA)-expressing nuclei represented a minor population, consistent with the reported proportion of fiber types^28^. Notably, *Myh2* expressing nuclei mainly located to the *Myh1*-positive side in the UMAP plot, but not to the *Myh4*-positive side (Fig. 1b). By fluorescence in situ hybridization (FISH), we could readily observe Myh1/Myh2 co-expressing fibers, but not Myh2/Myh4 fibers (Supplementary Fig. 3).

**Figure 1.**
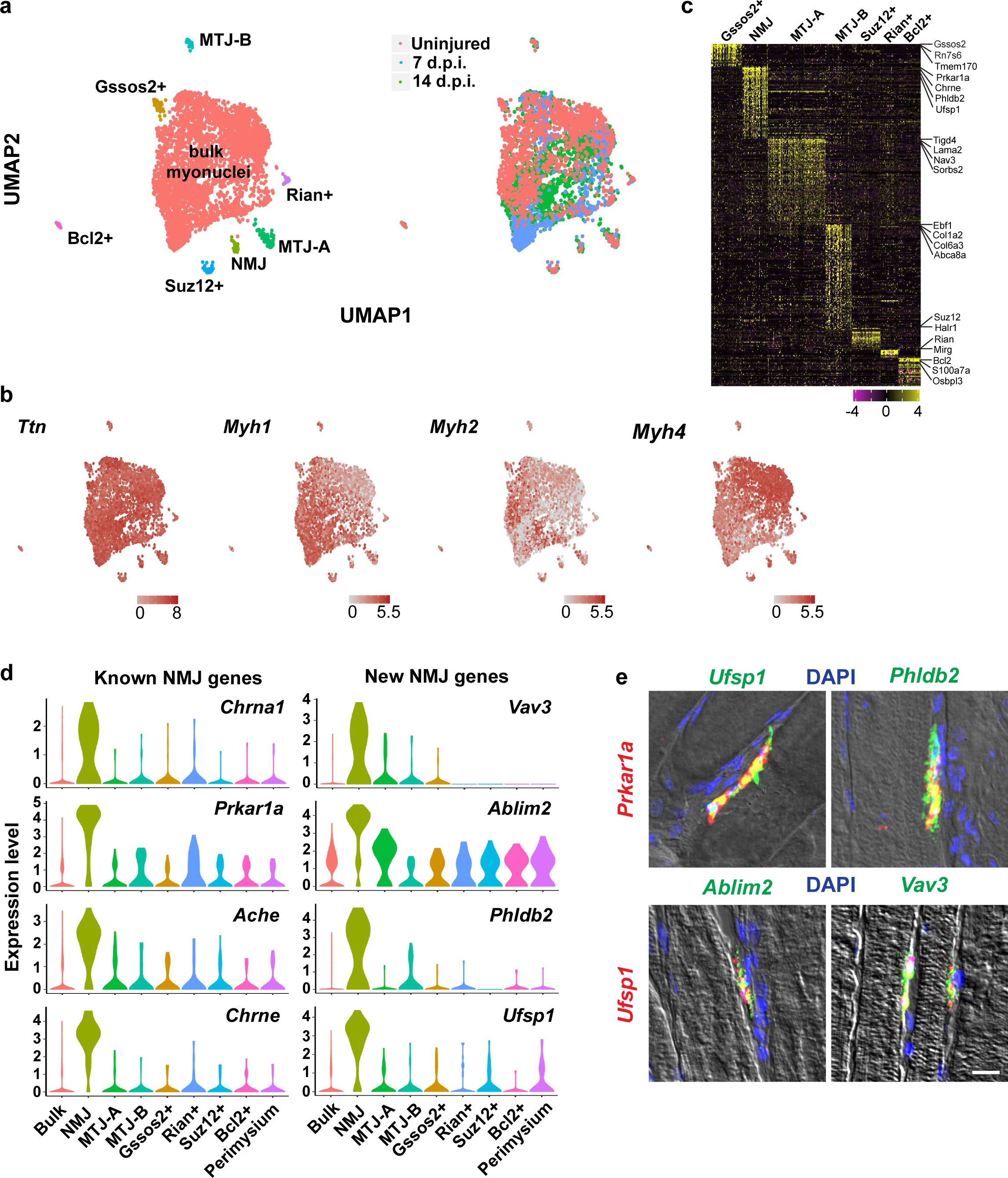
Nuclear heterogeneity in uninjured and regenerating muscles. **a**, UMAP plot of transcripts detected in nuclei of uninjured and regenerating muscles. The colors identify different nuclear populations (left) or nuclei in uninjured or regenerating muscle (right). **b**, Expression of *Ttn* and *Myosin* genes identifies myonuclei. **c**, Heat-map of specific genes enriched in clusters other than the bulk myonuclei. Top representative genes are indicated on the side. **d**, Violin plots of the previously known or newly identified NMJ marker genes. The perimysium population is identified after re-clustering the bulk myonuclei and described further in Figure 3. **e**, Upper row - conventional FISH against a known NMJ marker (*Prkar1a*) and newly identified NMJ genes (green). Bottom row – single molecule FISH against *Ufsp1* (red) and other newly identified NMJ genes (green). Expression patterns were validated in 2 or more individuals. Scale bar, 10 µm.

Next, we defined genes that showed a dynamic expression profile during regeneration (Supplementary Fig. 4a and Supplementary Table 1). For instance, genes like *Arrdc2, Smox, Gpt2* and *Pdk4* were strongly expressed in uninjured muscle but not during regeneration, whereas genes like *Mettl21c, Cish* and *Slc26a2* were specifically expressed at 7 d.p.i. These results were validated by RT-qPCR using isolated GFP+ myonuclei and FISH (Supplementary Fig. 4b-4c).

In addition to the bulk myonuclear population, we detected smaller populations with very distinct transcriptomes that expressed pan-muscle genes like *Ttn* (Fig. 1a-1c). Like the bulk nuclei, distinct nuclei in these populations expressed different myosin genes, indicating that the heterogeneity is not driven by fiber type differences (Fig. 1b). We first searched for and identified a cluster specifically expressing known NMJ marker genes such as *Chrna1, Prkar1a, Ache* and *Chrne* that was present in uninjured and regenerating muscle^5^ (Fig. 1d and Supplementary Table 4). Our data identified many other genes not previously known to be specifically expressed at the NMJ such as *Vav3, Ablim2, Phldb2* and *Ufsp1* (Fig. 1d). FISH of newly identified markers and the known marker *Prkar1a* confirmed their specific expression at the NMJ (Fig. 1e). The full list of marker genes identified in this study is available in Supplementary Table 2.

### Two distinct nuclear populations at the myotendinous junction

We found two clearly distinct nuclear populations that expressed MTJ-related genes in uninjured and regenerating muscle, and designated them MTJ-A and MTJ-B (Fig. 2a and Supplementary Table 4). MTJ-A nuclei expressed genes whose protein products are known to be enriched at the MTJ (e.g. *Itgb1*)^29^ as well as specific collagens (e.g. *Col24a1* and *Col22a1*). *Col22a1* has been functionally characterized in zebrafish using morpholino knockdowns that disrupt MTJ formation^30^. MTJ-B nuclei expressed an alternative set of collagens that are known to be deposited at the MTJ such as *Col1a2, Col6a1* and *Col6a3*^31^. *Col6a1* expression was particularly notable because its mutation causes Bethlem myopathy, which is characterized by deficits at the MTJ^32^.

**Figure 2.**
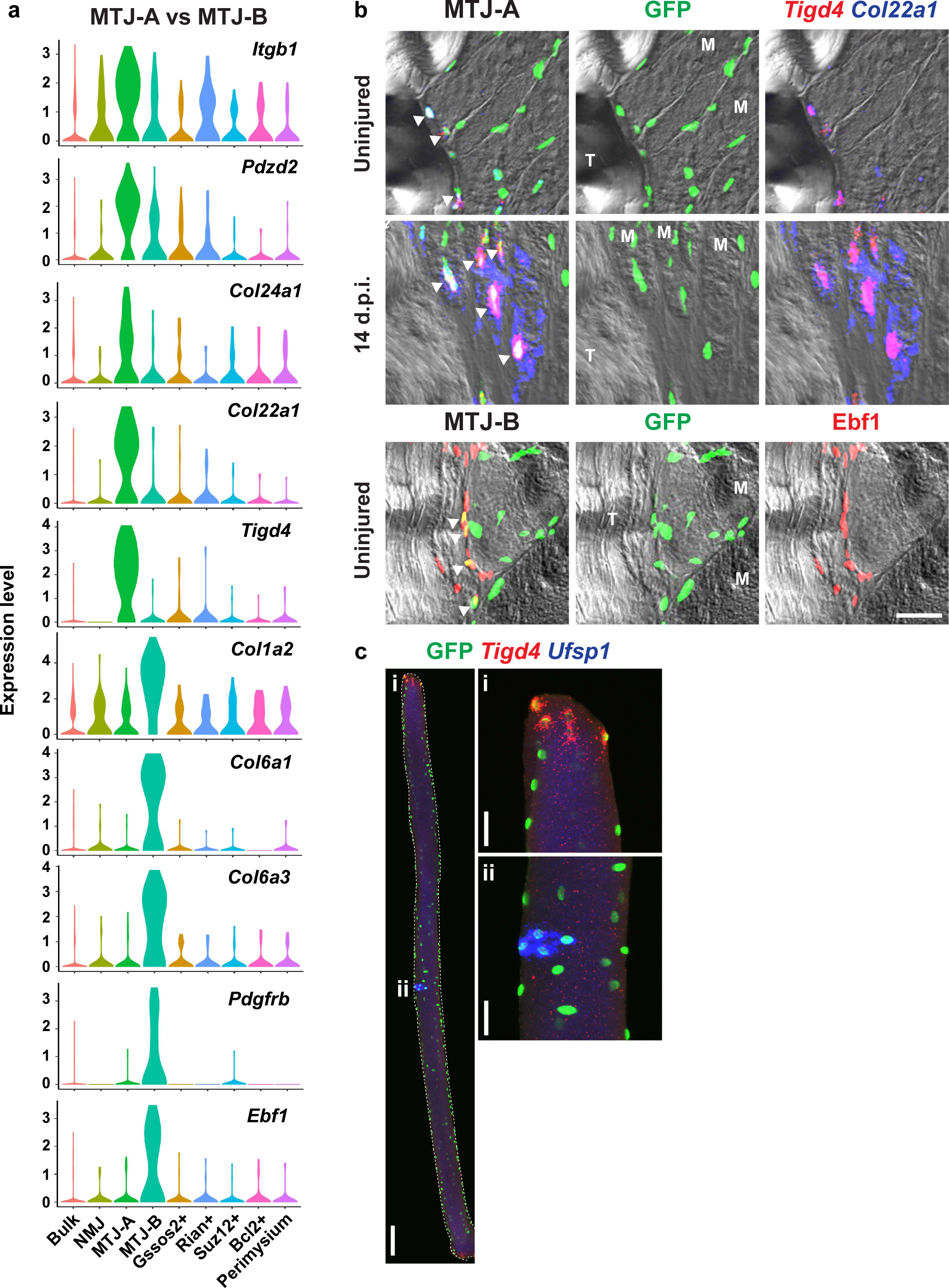
Two distinct nuclear populations at the myotendinous junction. **a**, Marker genes enriched in MTJ-A or MTJ-B nuclei are presented by violin plots. **b**, Upper two rows – single molecule FISH of two MTJ-A markers (*Tigd4* and *Col22a1*) in uninjured or 14 d.p.i TA muscle expressing H2B-GFP in myonuclei. Bottom row – Ebf1 (MTJ-B marker) immunoflourescence in uninjured TA muscle expressing H2B-GFP in myonuclei. Shown are MTJ regions; T, tendon and M, myofiber. Arrowheads indicate co-localization of MTJ marker genes and GFP. Scale bar, 30µm. **c**, Conventional FISH experiment in an isolated single EDL fiber. Insets show magnification of MTJ (i) and NMJ (ii) regions. Scale bars, 100 µm (for the entire fiber) and 30 µm (for the insets). Expression patterns were validated in 2 or more individuals.

We validated the two top marker genes of MTJ-A (*Tigd4* and *Col22a1*) using FISH and observed their expression in nuclei at fiber endings (Fig. 2b and Supplementary Fig. 5). These transcripts were exclusively expressed from H2B-GFP positive myonuclei and present only at the MTJ. Their expression became much more pronounced at 14 d.p.i. compared to uninjured muscle (Fig. 2b). To visualize heterogeneity within the syncytium, we isolated single fibers and performed double FISH (Fig. 2c). *Tigd4* FISH signals were detected at fiber ends where the MTJ is located, whereas *Ufsp1* transcripts appeared at the middle of the fiber where the NMJ is located.

We detected transcripts of MTJ-B genes (*Pdgfrb, Col6a3*) expressed from H2B-GFP nuclei at the MTJ in both uninjured muscle and at 14 d.p.i. (Supplementary Fig. 6). We also confirmed that Ebf1 protein is present in H2B-GFP positive myonuclei close to the MTJ (Fig. 2b). *Pdgfrb* and *Col6a3* are known to be expressed by the connective tissue, and indeed these transcripts were also detected in cells located distally to the MTJ and outside the fiber (Supplementary Fig. 6). However, such cells were neither marked by H2B-GFP nor by *Ttn*. Thus, MTJ-B nuclei co-expressing muscle genes and *Pdgfrb, Col6a3* or Ebf1 were exclusively located at end of the muscle fibers. Unlike MTJ-A nuclei, those expressing the MTJ-B signature were not found in every fiber. Because the MTJ-B signature includes both markers of muscle fibers and of connective tissue cells, we compared the gene signatures of MTJ-B nuclei to the ones of known cell types in the muscle tissue, specifically with the connective tissue cell types identified in a previous single-cell sequencing experiment that excluded syncytial myofibers^18^. None of these cell types expressed the MTJ-B signature. Thus, MTJ-B represents a novel nuclear population in the myofiber that co-expresses genes typical of the myofiber (e.g. *Ttn*) and of connective tissue (*Pdgfrb, Col6a3, Ebf1*).

### Identification of additional novel myonuclear populations

Further novel myonuclear compartments were identified by our systematic analysis, and we show exemplary genes preferentially expressed by each population in Figure 3a. The first of these we named after the top marker, *Rian*, a maternally imprinted lncRNA. This cluster also expressed other lncRNAs that are all located at the same genomic locus (also known as *Dlk1-Dio3* locus) like *Mirg* and *Meg3* (Fig. 3a-3b and Supplementary Table 2). The *Dlk1-Dio3* locus additionally encodes a large number of microRNAs expressed from the maternal allele, among them microRNAs known to target transcripts of mitochondrial proteins encoded by the nucleus^33^. FISH against *Rian* transcripts showed clear and strong expression in a subset of myonuclei (Fig. 3c), which was observed regardless of the animal’s sex (data not shown). FISH on isolated fibers showed dispersed localization of *Rian* expressing nuclei without clear positional preference (Fig. 3d). A previous study reported that the *Dlk1-Dio3* locus becomes inactive during myogenic differentiation^34^. However, our results show that some myonuclei retain expression, which might be important for the metabolic shaping of the fiber.

**Figure 3.**
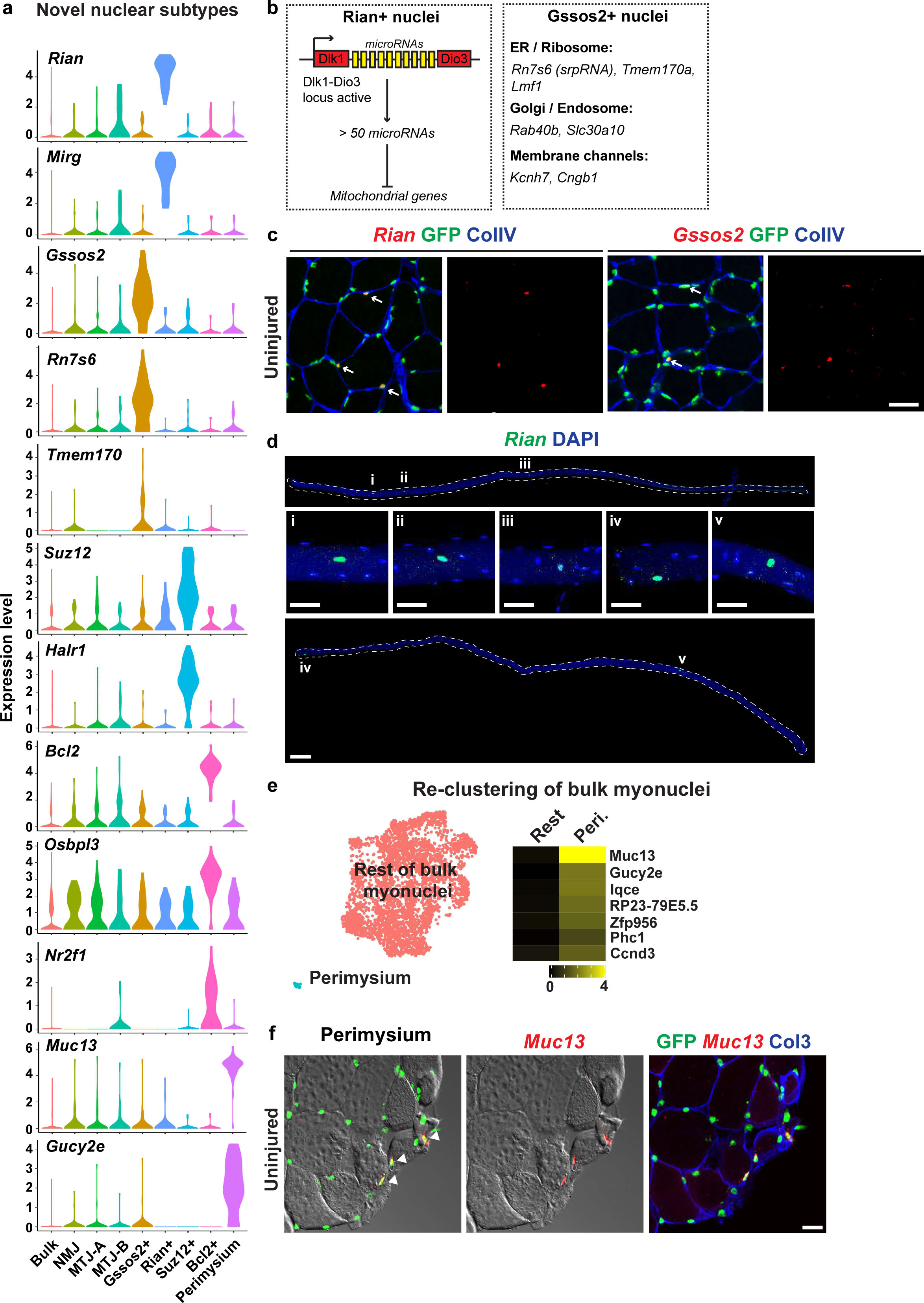
Identification of novel nuclear subtypes. **a**, Marker genes enriched in each of the novel nuclear population are presented by violin plots. **b**, Illustration of potential functions of Rian+ and Gssos2+ nuclei. Rian+ nuclei might regulate local mitochondrial metabolism through microRNAs embedded in *Dlk1-Dio3* locus, whereas Gssos2+ nuclei potentially regulate local protein synthesis and entry into the secretory pathway. **c**, Validation of the top markers of Rian+ and Gssos2+ nuclei by single molecule FISH in uninjured muscles. Note their strong expression in a subset of myonuclei (arrows). Scale bar, 30 µm. **d**, Expression of *Rian* in isolated EDL fibers. Insets show magnifications of indicated regions. Scale bars, 100 µm (for entire fibers) and 30 µm (for insets). **e**, (Left) UMAP plot of re-clustered bulk myonuclei identified in Figure 1a. (Right) Heat-map showing differentially expressed genes in perimysium (peri.) nuclei versus rest of the bulk myonuclei. Averaged gene expression levels are shown for each gene. **f**, Validation of *Muc13* expression in myonuclei adjacent to the perimysium by single molecule FISH in uninjured muscle. Scale bar, 30 µm. Expression patterns were validated in 2 or more individuals.

The top marker of the second cluster was *Gssos2*, an antisense lncRNA, and these nuclei expressed many genes that function in endoplasmic reticulum (ER)-associated protein translation and trafficking (Fig. 3a-3b). Among these were *Tmem170a* and *Rab40b*. Tmem170a induces formation of ER sheets, the site of active protein translation^35^, and Rab40b is known to localize to the Golgi/endosome and regulates trafficking^36^. Furthermore, the srpRNA (signal recognition particle RNA) *Rn7s6*, an integral component of ER-bound ribosomes, was markedly enriched in this population (Fig. 3a-3b). The enrichment of srpRNA was also observed when expression of repeat elements was quantified (Supplementary Fig. 7). FISH showed that *Gssos2* displayed a heterogeneous and strong expression in a subset of myonuclei and that *Rian* and *Gssos2* were expressed in different nuclei; we examined more than 100 Rian+ or Gssos2+ nuclei from 2 individuals and did not observe co-expressing nuclei. (Fig. 3c; Supplementary Fig. 8a). *Rian* and *Gssos2* were located away from NMJ nuclei (Supplementary Fig. 8b). Therefore, Rian+ and Gssos2+ nuclei represent independent nuclear populations.

Two remaining populations (Suz12+ and Bcl2+ nuclei) need further characterization. The top two markers expressed by Suz12+ nuclei were *Suz12*, a core Polycomb complex component, and *Halr1* transcripts, a long non-coding RNA expres sed from the *Hoxa* locus, suggesting that specific mechanisms of epigenomic regulation might be used in these nuclei. Bcl2+ nuclei strongly expressed genes involved in steroid signaling such as *Osbpl3* (oxysterol-binding protein) and *Nr2f1* (steroids-sensing nuclear receptor).

Re-clustering of the bulk myonuclei in Figure 1a revealed an additional nuclear subpopulation (Fig. 3e and Supplementary Fig. 9). This new subpopulation was characterized by the enrichment of marker genes such as *Muc13* and *Gucy2e* (Fig. 3a and 3e). FISH showed that myonuclei expressing *Muc13* were always located at the very outer part of the muscle tissue near the perimysium (Fig. 3f). A previous ultrastructural study suggested that myofibers and perimysium establish specialized adhesion structures^37^, and our data suggest that we have detected a myonuclear compartment participating in this process.

### snRNAseq of fibers in *Mdx* dystrophy model

To begin to understand whether and how myonuclear heterogeneity is altered in muscle disease, we conducted snRNAseq on *Mdx* fibers (1,939 nuclei), a mouse model of muscular dystrophy caused by mutation of the *Dystrophin* gene (Fig. 4a and Supplementary Fig. 10a). To examine how the transcriptome of *Mdx* myonuclei is related to those of uninjured and regenerating muscle, we calculated gene signature scores of each nucleus based on the top 25 genes that distinguish uninjured and regenerating fibers. This showed that nuclei of the *Mdx* muscle mostly resembled those from the uninjured and 14 d.p.i muscle, whereas the signature of 7 d.p.i myofibers was depleted (Fig. 4b). Cluster A displayed marker genes that were largely shared with those specific to uninjured fibers like *Arrdc2, Glul, Smox* and *Gpt2* (Fig. 4c and Supplementary Fig. 4a) and might correspond to nuclei from fibers that are little damaged or undamaged.

**Figure 4.**
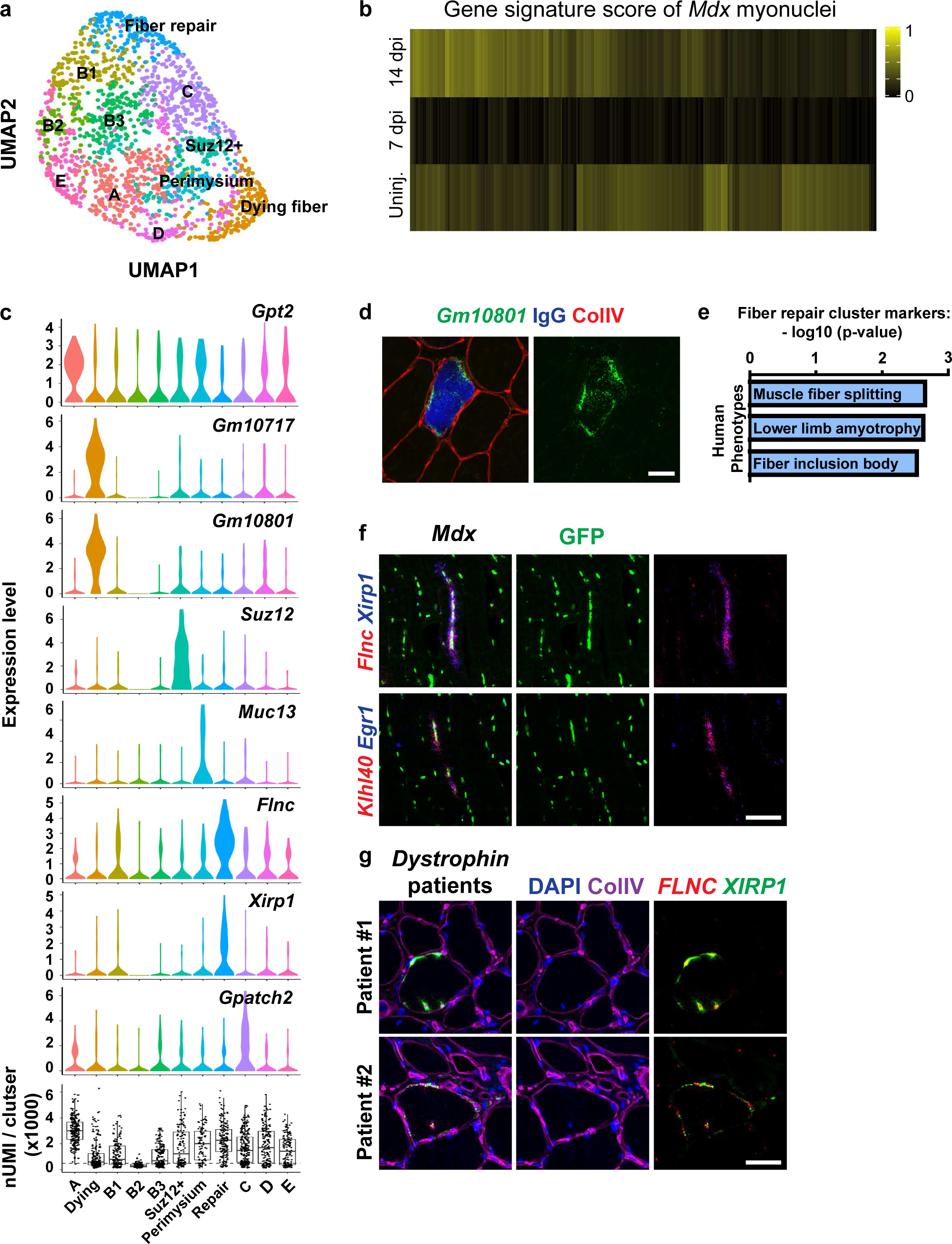
snRNAseq analysis of *Mdx* muscle. **a**, UMAP plot of *Mdx* myonuclei (1,939 nuclei). **b**, Each *Mdx* myonucleus was assigned a gene signature score, i.e. a gene expression score indicating similarity with uninjured (uninj.), 7 or 14 d.p.i. myonuclei. Each column represents an individual nucleus. **c**, Marker genes enriched in each population are presented by violin plots. The bottommost histogram shows nUMI in each population. **d**, The ncRNA *Gm10801* is expressed in IgG-positive fibers, indicating that it defines nuclei of necrotic fibers. **e**, Fiber repair myonuclei express high levels of various genes implicated in fiber repair that are implicated in myopathies. The p-values were corrected using the Benjamini-Hochberg (BH) procedure. **f**, Indicated marker genes of the cluster in e are co-expressed in longitudinal muscle sections of *Mdx* mice. **g**, Co-expression of *FLNC* and *XIRP1* in the muscle from dystrophy patients carrying mutations in the *DYSTROPHIN* gene (*DYS* del exons15-18; *DYS* c.2323A>C). Control images for f and g are shown in Supplementary Fig. 11a. All scale bars, 50 µm. Expression patterns were validated in 2 or more individuals.

We found novel nuclear populations in the *Mdx* dataset that were not identified in the uninjured/regenerating muscle. The first of these highly expressed various non-coding transcripts, and further experiments demonstrated that nuclei expressing these transcripts were located inside dying fibers (Fig. 4c and Supplementary Fig. 10b). In particular, staining with mouse IgG that identifies fibers with leaky membranes demonstrated these ncRNAs were expressed in IgG+ fibers (Fig. 4d). Further, fibers strongly expressing these ncRNAs were highly infiltrated with H2B-GFP negative cells likely corresponding to macrophages (Supplementary Fig. 10c-10d). In line with the idea that these nuclei represent dying fibers, they had low UMI counts suggesting low transcriptional activity or high mRNA degradation (bottom histogram in Fig. 4c). Whether the ncRNAs are the consequence or active contributors to fiber death needs further investigation. In addition, three populations (B1-B3) located adjacent to each other in the UMAP map and had low UMI counts, but did not display any clear marker genes. We speculate that these nuclei might also originate from damaged fibers.

Next, we searched for the clusters identified in the uninjured and regenerating muscle. In the UMAP of the *Mdx* dataset, clusters corresponding to NMJ, MTJ-A, MTJ-B, Rian+, Gssos2+ and Bcl2+ populations were not identifiable in the UMAP. However, we detected two subpopulations also present in uninjured/regenerating muscle, Suz12+ and perimysial nuclei, pointing to some degree of specificity (Fig. 4a). We therefore used gene signature scores to identify nuclei that display a correlative expression of signature genes. Such inspection showed that MTJ-A nuclei were present in the UMAP but did not cluster together (Supplementary Fig. 10e). Nevertheless, marker genes of MTJ-A (*Tigd4* and *Col22a1*) robustly labeled the MTJ of *Mdx* muscle (Supplementary Fig. 10f). We speculate that fiber-level heterogeneity (e.g. dying fibers, intact fibers and regenerating fibers) drives the shape of the UMAP map in *Mdx*, which might interfere with the clustering of MTJ-A nuclei. Unlike MTJ-A, nuclei with high signature scores of NMJ, MTJ-B, Rian+, Gssos2+ and Bcl2+ nuclei were not detected. We investigated the expression of NMJ genes in further depth. This showed that the strict co-expression of two NMJ marker genes (*Ufsp1* and *Prkar1a*) typical for the control muscle was lost, and that these genes were instead expressed in a dispersed manner in the *Mdx* muscle (Supplementary Fig. 10f). Notably, the histological structure of the NMJ is known to be fragmented in *Mdx* mice^38,39^, and our data suggest that also postsynaptic nuclei are incompletely specified.

### Emergence of a nuclear population implicated in fiber damage repair in *Mdx* model

Marker genes of the cluster ‘fiber repair’ in Figure 4a showed enrichment of ontology terms related to human muscle disease (Fig. 4e). Indeed, many top marker genes were previously reported to be mutated in human myopathies (*Flnc, Klhl40*, and *Fhl1*)^40-42^ or to directly interact with proteins whose mutation causes disease (Ahnak interacts with dysferlin; Hsp7b or Xirp1 interact with Flnc)^43-45^. Combinatorial FISH in tissue sections confirmed co-expression of such marker genes in a subset of nuclei of *Mdx* muscle, but such nuclei were not present in control muscle (Fig. 4f, Supplementary Fig. 11a-11b). Further, nuclei expressing these genes were frequently closely spaced in fibers. We also verified that *FLNC* and *XIRP1* were co-expressed in nuclei from patient biopsies with confirmed *DYSTROPHIN* mutation, but not in healthy human muscle (Fig. 4g and Supplementary Fig. 11a).

Previous studies have established that Flnc and Xirp1 proteins localize to sites of myofibrillar damage to repair such insults^44,46,47^, whereas Dysferlin, an interaction partner of Ahnak, functions during repair of muscle membrane damage^48^. Our analysis shows that these genes are transcriptionally co-regulated which might occur in response to micro-damage. To substantiate that this signature is not specific to muscular dystrophy caused by *Dystrophin* mutation, we investigated whether they can be identified in *Dysferlin* deficient muscle where the continuous micro-damage to the membrane is no longer efficiently repaired^48^. Again, we observed nuclei co-expressing *Flnc* and *Xirp1* in this mouse disease model and in biopsies from human patients with *DYSFERLIN* mutations (Supplementary Fig. 11c-11d). We propose that the genes that mark this cluster represent a ‘repair’ signature. Notably, the accompanying paper (Petrany et al.) identified a similar population during late postnatal development and in the aging muscle, indicating that the ‘repair’ genes might also function during fiber remodeling.

Finally, we identified another new population in the *Mdx* muscle, cluster C, that expressed marker genes such as *Gpatch2, Emilin1* and *Pde6a* not previously studied in a muscle context (Fig. 4c and Supplementary Table 2). The role of this population in muscle pathophysiology needs further characterization.

### Nuclear heterogeneity in muscle spindle fibers

In principle, our approach can be used to explore nuclear heterogeneity in specific fiber types. We thus aimed to investigate heterogeneity in muscle spindles that detect muscle stretch and function in motor coordination^49^. Muscle spindles are extremely rare and ∼10 spindles exist in a TA muscle of the mouse^50^. They contain bag and chain fibers, and their histology suggests further compartmentalization (Fig. 5a). *HSA-Cre* labels myonuclei of the spindle (Fig. 5b), but the overwhelming number of nuclei derive from extrafusal fibers. To overcome this, we used *Calb1-Cre* to specifically isolate spindle myofiber nuclei (Fig. 5b) and discovered different nuclear subtypes inside these specialized fibers (Fig. 5c).

**Figure 5.**
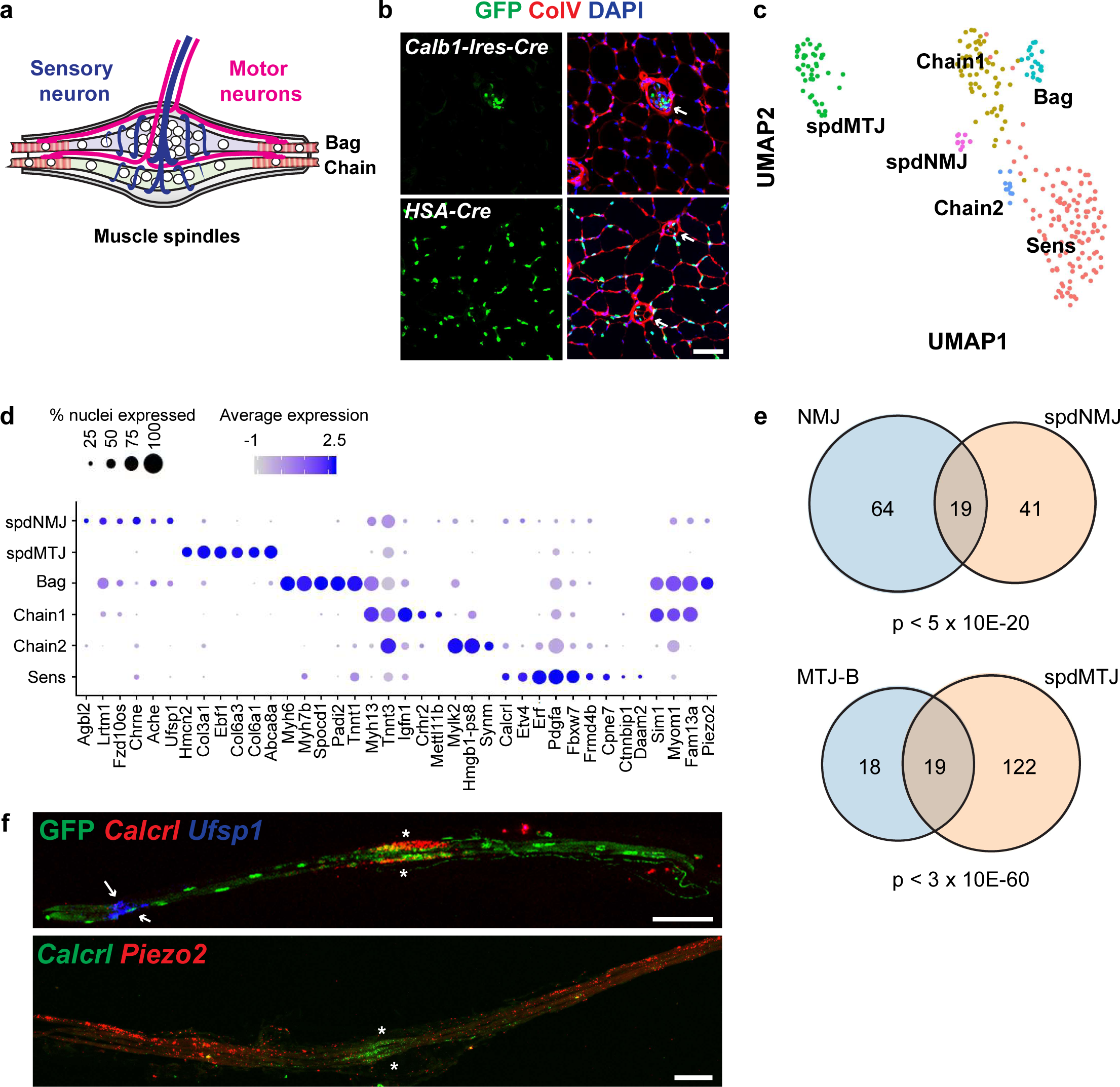
Functional compartments inside muscle spindle fibers. **a**, Schema showing the structure of the muscle spindle. **b**, Specific labeling of spindle myonuclei using *Calb1-Ires-Cre*. Arrows indicate muscle spindles. **c**, UMAP plot of muscle spindle myonuclei (260 nuclei). **d**, Expression map of nuclear populations identified in c. **e**, Venn diagram comparing ‘spdNMJ vs extrafusal fiber NMJ’ and ‘spdMTJ vs extrafusal fiber MTJ-B’. Genes enriched in each population (average logFC > 0.7) were used to generate the diagrams. Statistical analysis was performed using hypergeometric test using all genes detected in uninjured/regenerating and spindle datasets as background. **f**, Single molecule FISH experiments of isolated muscle spindle fibers. Arrows indicate spdNMJ, and asterisks the central non-contractile parts of spindle fibers. All scale bars, 50 µm. Expression patterns were validated in 2 or more individuals.

Bag fibers are slow fibers and express *Myh7b*^51,52^, whereas chain fibers are fast. A cluster expressing *Myh7b* and *Tnnt1*, a slow type troponin isoform, was assigned to mark Bag fibers. In contrast, two clusters expressed *Myh13*, a fast type Myosin, or *Tnnt3*, a fast type troponin, which we named Chain1 and Chain2. Strikingly, we identified a cluster that expressed a set of genes largely overlapping with those identified in NMJ nuclei of extrafusal fibers, e.g. *Chrne, Ufsp1* and *Ache*, which we assign as the NMJ of the spindle (spdNMJ) (Fig. 5d and 5e). Furthermore, the spindle myotendinous nuclei (spdMTJ) expressed a significantly overlapping set of genes as those identified in MTJ-B nuclei of extrafusal fibers (Fig. 5d and 5e). MTJ-A markers were not detected. Notably, the clusters Bag and spdNMJ expressed the mechanosensory channel *Piezo2*^53^. To verify the assignment and to define the identity of an additional large compartment (labeled as Sens), we validated the expression of different marker genes in H2B-GFP positive fibers of *Calb1-Cre* muscle in tissue sections (Supplementary Fig. 12b) and in fibers after manual isolation (Fig. 5f). FISH of *Calcrl*, a marker of the Sens cluster, showed specific localization to the central part of spindle fibers containing densely packed nuclei, the site where sensory neurons innervate (Fig. 5f). In the same fiber, transcripts of the spdNMJ marker gene *Ufsp1* located laterally as a distinct focus. In contrast, *Piezo2* was expressed throughout the lateral contractile part of the fiber, but was excluded from the central portion. Thus, the central non-contractile part of the muscle spindle that is contacted by sensory neurons represents a fiber compartment with specialized myonuclei clearly distinguishable from the spdNMJ.

### Profiling transcriptional regulators across distinct compartments

To gain insights into the transcriptional control of the different nuclear compartments, we investigated the expression profile of transcription factors and epigenetic regulators (Fig. 6a and Supplementary Table 3). Notably, the transcript encoding Etv5 (also known as Erm), a transcription factor known to induce the NMJ transcriptome^54^, and its functional homolog Etv4 were enriched in NMJ nuclei. Irf8 (3rd rank factor in NMJ) is also interesting as mutation of an Irf8 binding site in the *CHRNA* promoter causes *CHRNA* misexpression in the thymus and leads to the autoimmune disease myasthenia gravis^55^, implicating Irf8 in the control of an NMJ gene in a tissue outside of the muscle. In MTJ-A nuclei, Smad3, the effector of TGF-β signaling, was found as the second rank factor. In addition, TGF-β receptors were also expressed by MTJ-A myonuclei. TGF-β is released by force from tenocytes and is required to maintain tenocytes^56^, but our dataset suggests that MTJ myonuclei can also receive TGF-β signals.

**Figure 6.**
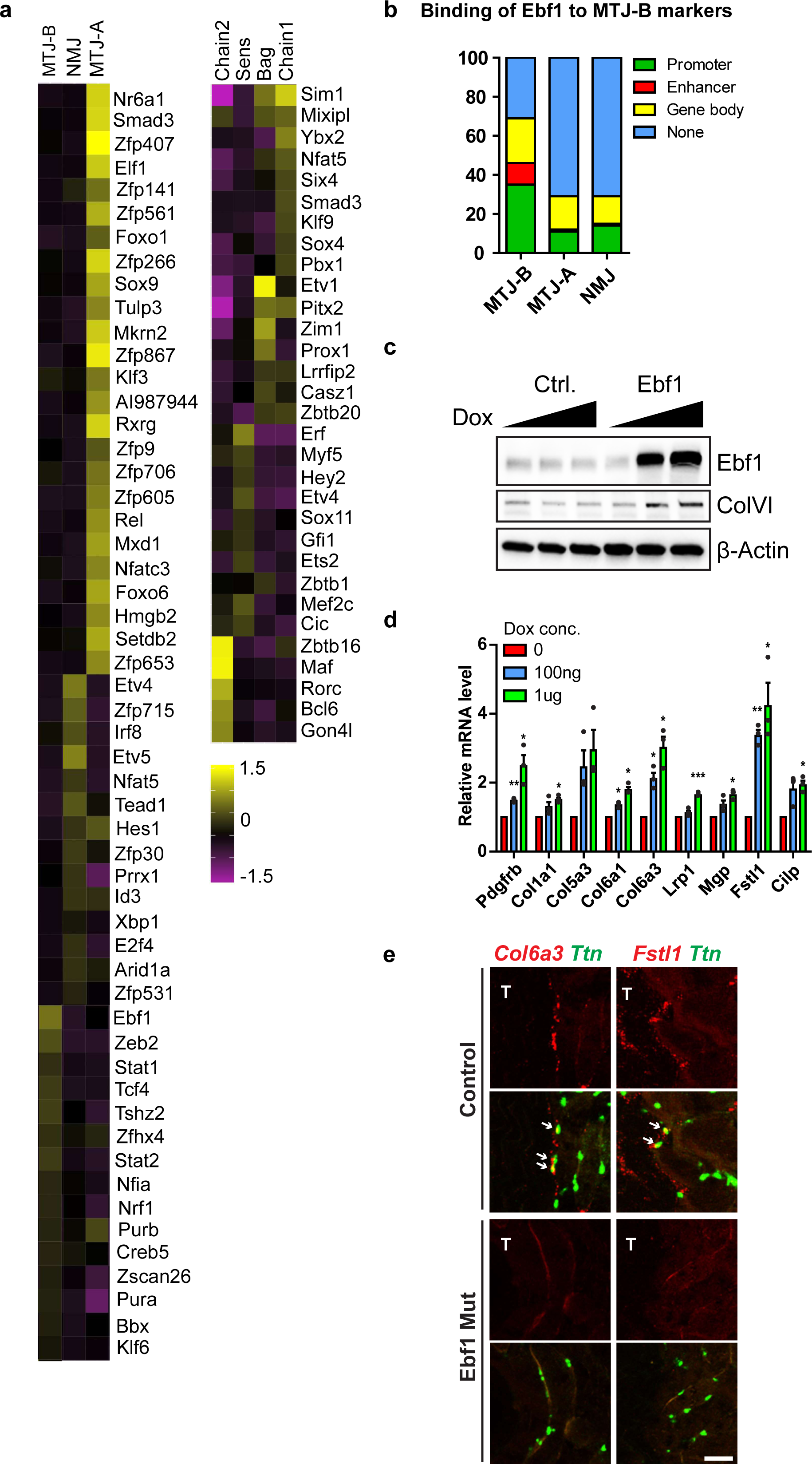
Expression profiles of transcription factors and epigenetic regulators across distinct nuclear subtypes. **a**, Heat-map showing the expression level of transcription factors and epigenetic regulators in indicated nuclear subtypes; see also Supplementary Table 3 for the full list including nuclear subtypes not shown in this Figure. **b**, Ebf1 directly binds to genomic regions of the top 100 genes identified to be specifically expressed in MTJ-B nuclei (ENCODE project - accession number ENCSR000DZQ), but much less to markers of MTJ-A or NMJ. Classification of the Ebf1 binding sites in these 100 genes. **c**, Western blot analysis of C2C12 cell line expressing doxycycline-inducible Ebf1. **d**, Indicated genes were analyzed by RT-qPCR before/after inducing Ebf1 expression. Dox treatment in control cells (no Ebf1) did not have any effect (data not shown). Error bars indicate S.E.M. Two tailed paired student’s t-test between untreated and Dox treated cells (n=3). *, p < 0.05. **, p < 0.01. ***, p < 0.001. **e**, Single molecule FISH of indicated marker genes in TA muscle of control or *Ebf1* mutant mice. Arrows indicate myonuclei expressing MTJ-B marker genes. T, tendon. Downregulation of expression was validated in 4 individuals. Scale bar, 50 µm.

To test whether our dataset can identify a novel and functionally relevant factor, we chose to further study Ebf1, the most strongly enriched transcription activator in MTJ-B. ChIP-Seq data of Ebf1 (ENCODE project - accession number ENCSR000DZQ) showed that Ebf1 directly binds to ∼70% of the genes we identified as MTJ-B markers (Fig. 6b). In contrast, Ebf1 binds less than 30% of NMJ or MTJ-A marker genes. We generated a C2C12 cell line in which Ebf1 expression was induced by doxycycline (Fig. 6c). RT-qPCR analysis of selected markers showed that many of them were induced in a dose-dependent manner upon Ebf1 over-expression (Fig. 6d), as was ColVI protein (Fig. 6c). To validate these findings *in vivo*, we analyzed *Ebf1* mutant muscle. Using FISH for *Ttn* to identify myonuclei, we observed a strong reduction of *Col6a3* and *Fst1l* transcripts in the *Ebf1* mutant compared to control MTJ myonuclei (Fig. 6e). Their expression from non-myonuclei that also express Ebf1 was also diminished. In contrast, the expression of NMJ, MTJ-A and *Rian* markers was not affected in *Ebf1* mutants (Supplementary Fig. 13). Taken together, our dataset provides a template for the identification of regulatory factors that establish or maintain these compartments.

## Discussion

Here, we used snRNAseq to systemically characterize the transcriptional heterogeneity of myofiber nuclei. Common to all was the expression of muscle-specific genes like *Ttn*, but small subpopulations were detected that expressed an additional layer of distinct and characteristic genes. Our analysis of the uninjured and regenerating muscle identified nuclear populations at anatomically distinct locations such as the NMJ and MTJ-A populations that were known to exist, as well as MTJ-B and perimysial nuclei, two populations that are first described here. Moreover, we found a number of new populations that are scattered throughout the myofiber, among them the two distinct Rian*+* and Gssos2*+* nuclear subtypes. Thus, myonuclear populations are not always associated with distinctive anatomical features. How these nuclear subtypes emerge, i.e. whether they are associated with other cell types in the muscle tissue or arise stochastically needs further study. Collectively, our data identified many genes that are specifically expressed in the various nuclear populations, providing a comprehensive resource for studying these compartments. We provide a webserver where users can freely explore the expression profile of their gene of interest in myonuclei (https://shiny.mdc-berlin.net/MyoExplorer/). Together, our results reveal the complexity of the regulation of gene expression in the syncytium and show how regional transcription shapes the architecture of multinucleated skeletal muscle cells.

Our analysis also shows that the transcriptional heterogeneity in myonuclei is dynamic. For instance, during regeneration the gene expression signatures of bulk nuclei differ from those of the uninjured muscle, and differences between early and later stages of regeneration can be detected. Further, the frequency of different myonuclear subtypes might suggest dynamic changes in nuclear compartments during regeneration (Supplementary Table 4). However, the proportion of these nuclear subtypes is low (0.5∼3%) which precludes a definitive conclusion at this stage. Finally, FISH experiments showed increased expression of MTJ-A marker genes during regeneration, indicating that a higher demand for the products of MTJ genes might exist when the MTJ needs to be re-established, whereas less are needed to maintain this structure.

Another dynamic aspect of myonuclear heterogeneity is demonstrated by our analysis on dystrophic muscles. snRNAseq of *Mdx* muscles revealed a number of novel compartments. In particular, we identified the molecular signature of degenerating fibers and a transcriptional program that appears to be associated with fiber repair. These newly identified gene signatures might be useful for a quantitative and rapid assessment of muscle damage in the clinic. In addition to the appearance of new compartments, many nuclear subtypes present in normal muscle were lost in the *Mdx* muscle, including nuclei expressing the NMJ signature. The protein (but not the transcript) encoded by *Dystrophin* is known to be highly enriched at the NMJ, and the NMJ was previously observed to be functionally abnormal in *Mdx* mice^57-59^. The absence of nuclei that express the NMJ signature in the *Mdx* muscle might provide a molecular correlate for these deficits. Together, our analysis demonstrates that the use of snRNAseq can provide novel insights into the molecular pathophysiology of muscle disease.

Given the large size and complexity of the muscle tissue, the full diversity of myonuclei likely needs further exploration. Here, we concentrated our analysis to a single muscle group, the tibialis anterior muscle that mainly contains fast fibers, and determined the transcriptional heterogeneity and programs in uninjured, regenerating and dystrophic muscle. An accompanying manuscript (Petrany et al.) successfully used snRNAseq to define nuclear subtypes in the postnatal, adult and aged tibialis anterior muscle. The two studies identified an overlapping set of compartments, but each also found distinct ones, underlining the fact that transcriptional compartments in the muscle are highly dependent on variables like disease or age. Further, different isolation strategies and snRNA sequencing methods were used in the two studies. Our strategy identified the rare MTJ-B or perimysial nuclei subtypes that were not detected by others, and thus the strategy of genetic labeling and isolating myonuclei should be promising for the identification of new nuclear subtypes in other muscle groups and contexts. Nevertheless, nuclei from the muscle spindle, a very rare and specialized fiber type, were not detected in any of the datasets that analyzed a random set of myonuclei, but we overcame this limitation by restricting the genetic labeling. The snRNAseq analysis of spindle nuclei revealed many subtypes inside these rare fibers, especially the presence of a specific compartment at the site of innervation by proprioceptive sensory neurons. More generally, our approach should be useful to investigate other syncytial cell types such as the placental trophoblasts or osteoclasts.

## Methods

### Isolation of nuclei from TA muscle

For each sorting of uninjured, regenerating or *Mdx* muscles, we pooled two TA muscles from two mice (one TA from each mouse). Dissected TA muscles were minced into small pieces in a 3.5 cm dish on ice with scissors in 300 µl hypotonic buffer (250 mM sucrose, 10 mM KCl, 5 mM MgCl2, 10 mM Tris-HCl pH 8.0, 25mM HEPES pH 8.0, 0.2 mM PMSF and 0.1 mM DTT supplemented with protease inhibitor tablet from Roche), and transferred with a 1 ml pipette and cut tip to a 2 ml ‘Tissue homogenizing CKMix’ tube (Bertin instruments KT03961-1-009.2) containing ceramic beads of mixed size. The dishes and tips were washed with 700 µl hypotonic buffer. Samples were incubated on ice for 15 minutes and homogenized with Precellys 24 tissue homogenizer (Bertin instruments) for 20 seconds at 5,000 rpm. Homogenized samples were passed once through 70 µm filter (Sysmex), twice through 20 µm filter (Sysmex), and once though 5 ml filter cap FACS tube (Corning 352235). DAPI (Sigma) was added to final concentration of 300 nM to label DNA. GFP and DAPI double positive nuclei were sorted using ARIA Sorter III (BD).

For isolation of muscle spindle nuclei, we employed two different protocols. In the first protocol (protocol 1 in Supplementary Fig. 12a), we used 6 TA muscles from 3 mice (3 TA muscles per 2 ml homogenizer tube) and used the same procedure as described above. From this, we isolated 96 nuclei. For the second procedure (protocol 2 in Supplementary Fig. 12a), we used 8 TA muscle from 4 mice (4 TA muscles per 2 ml homogenizer tube) and aimed to shorten the isolation time. For this, 0.1% Triton X-100 was added to the hypotonic buffer to solubilize the tissue debris, which are otherwise detected as independent particles during FACS. After homogenization and filtration nuclei were pelleted by centrifuging at 200 g for 10 minutes at 4°C. After aspirating the supernatant, the pellet was resuspended in 300 µl hypotonic buffer (without detergent) and passed through the FACS tube. The subsequent FACS sorting yielded 192 spindle nuclei.

### Library generation and sequencing

96-well plates for sorting were prepared using an automated pipetting system (Integra Viaflo). Each well contained 1.2 µl of master mix (13.2 µl 10% Triton X-100, 25 mM dNTP 22 µl, ERCC spike-in 5.5 µl and ultrapure water up to 550 µl total) and 25 ng/µl barcode primers. Plates were stored at -80°C until use.

After sorting single nuclei into the wells, plates were centrifuged at 4,000 g for 1 minute, incubated on incubated on 65°C for 5 minutes and immediately cooled on ice. Subsequent library generation was performed using the CEL-Seq2 protocol as described^26^. After reverse transcription and second strand synthesis, products of one plate were pooled into one tube and cleaned up using AMPure XP beads (Beckman Coulter). After in vitro transcription and fragmentation, aRNA was cleaned up using RNAClean XP beads (Beckman Coulter) and eluted in 7 μl ultrapure water. 1 μl of aRNA was analyzed on Bioanalyzer RNA pico chip for a quality check. To construct sequencing library, 5 μg aRNA was used for reverse transcription (Superscript II, Thermofisher) and library PCR (Phusion DNA polymerase, Thermofisher). After clean up using AMPure XP beads, 1 μl sample was ran on Bioanalyzer using a high sensitivity DNA chip to measure size distribution, which we demonstrated the presence of a peak of around 400 bp length. Further information on sequencing platforms and multiplexing are available in Supplementary Table 5.

### Bioinformatics analysis

Single nucleus RNA-Sequencing data was processed using PiGx-scRNAseq pipeline - a derivative of a CellRanger pipeline, but enabling deterministic analysis reproducibility (version 0.1.5)^60^. In short, polyA sequences were removed from the reads. The reads were mapped to the genome using STAR^61^. Number of nuclei, for each sample, was determined using dropbead^62^. Finally, a combined digital expression matrix was constructed, containing all sequenced experiments.

Digital expression matrix post processing was performed using Seurat^63^. The raw data was normalized using the NormalizeData function. The expression of each nucleus was then normalized by multiplying each gene with the following scaling factor: 10000/(total number of raw counts), log(2) transformed, and subsequently scaled. Number of detected genes per nucleus was regressed out during the scaling procedure.

Variable genes were defined using the FindVariableGenes function with the default parameters. Samples were processed in three groups with differing parameters. Samples originating from uninjured, 7 d.p.i. and 14 d.p.i. were processed as one group, samples from the *Mdx* mouse as the second group and muscle spindle nuclei as the third group. The samples originating from different biological sources contained markedly different properties – number of detected genes and UMIs, which precluded their analysis with the same parameter set.

To test stress response in our dataset, we used the signature of stress-induced genes identified previously^23^. We tested whether the stress-induced genes were co-expressed in individual cells using two different algorithms, AddModuleScore from Seurat and AUCell. The distribution of obtained scores was similar regardless of the algorithms used. Based on this, we concluded that there were only a handful nuclei (less than 10 among all the nuclei analyzed in total) which showed co-expression of known stress genes, and could therefore be considered “stressed”.

For samples of the first group, nuclei with less than 500 detected genes were filtered out. Subsequently, genes which were detected at least in 5 nuclei were kept for further analysis. To remove the putative confounding effect between time of sample preparation and biological variable (injury), the processed expression matrices were integrated using the FindIntegrationAnchors function with reciprocal PCA, from the Seurat package. The function uses within batch covariance structure to align multiple datasets,

The integration was based on 2000 top variable features, and first 30 principal components. UMAP was based on the first 15 principal components. Outlier cluster detection was done with dbscan (10.18637/jss.v091.i01.), with the following parameters eps = 3.4, minPts = 20.

*Mdx* samples contain dying fibers, which have few detected transcripts, as well as others that resemble uninjured fibers. Thus, *Mdx* nuclei show a big variance in the number of detected genes. Therefore, *Mdx* samples were processed by filtering out all nuclei with less than 100 detected genes. Top 100 most variable genes were used for the principal component analysis. UMAP and Louvain clustering were based on the first 15 principal components. Resolution parameter of 1 was used for the Louvain clustering.

In muscle spindle nuclei, we detected fewer genes than in uninjured fibers, but the variance was low. Spindle cell samples were processed by filtering out all cells with less than 300 detected genes. The top 200 most variable genes were used for the principal component analysis. UMAP and Louvain clustering were based on the first 15 principal components. Resolution parameter of 1 was used for the Louvain clustering.

For all datasets, multiple parameter sets were tested during the analysis, and the choice of parameters did not have a strong influence on the results and the derivative biological conclusions. Genes with cluster specific expression were defined using Wilcox test, as implemented in the FindAllMarkers function from the Seurat package. Genes that were detected in at least 25% of the cells in each cluster were selected for differential gene expression analysis.

NMJ Nuclei Definition: NMJ nuclei were identified based on the expression of three previously known markers (*Prkar1a, Chrne* and *Ache*), using the AddModuleScore function from the Seurat package. All cells with a score greater than 1 were selected as NMJ positive cells. NMJ marker set was expanded by comparing the fold change of gene expression in averaged NMJ positive to NMJ negative cells.

MTJ A and B Nuclei Definition: The original gene sets were extracted from cluster specific genes detected in the uninjured, 7 d.p.i. and 14 d.p.i. experiment. Cells were scored as MTJ A/B using the aforementioned gene set, with the AddModuleScore function. All cells which had a respective score greater than 1 were labeled as MTJ A/B positive cells.

*Mdx* nuclei scoring by uninjured and regenerating signatures: First, gene signatures specific for each time point were selected using the FindAllMarkers function from the Seurat library, using the default parameters. *Mdx* samples were scored using the top 25 genes per time point with the AUCell method (10.1038/nmeth.4463).

Repetitive element annotation was downloaded from the UCSC Browser database (10.1093/nar/gky1095) on 21.01.2020. Pseudo-bulk bigWig tracks were constructed for each cluster in uninjured, 7.d.p.i, and 14.d.p.i. The tracks were normalized to the total number of reads. Repetitive element expression was quantified using the ScoreMatrixBin function from the genomation (10.1093/bioinformatics/btu775) package, which calculates the average per-base expression value per repetitive element. The expression was finally summarized to the repetitive element family (class) level by calculating the average expression of all repeats belonging to the corresponding family (class).

Transcription factor compendium, used in all analyses was downloaded from AnimalTFDB (https://doi.org/10.1093/nar/gky822). The expression map (Fig. 5d) for spindle fibers was created using the DotPlot from the Seurat package on a selected set of cluster specific genes. Gene ontology analysis was performed using the Enrichr program (10.1093/nar/gkw377). (http://amp.pharm.mssm.edu/Enrichr/).

The online tool for interactive exploration of the single-cell data – MyoExplorer, was set up using iSEE (10.12688/f1000research.14966.1).

### Mouse lines and muscle injury

All experiments were conducted according to regulations established by the Max Delbrück Centre for Molecular Medicine and LAGeSo (Landesamt für Gesundheit und Soziales), Berlin. *HSA-Cre, Calb1-IRES-Cre* and *Mdx* mice were obtained from the Jackson laboratory. *Rosa26-Lsl-H2B-GFP* reporter line was a kind gift from Martin Goulding (Salk institute) and was described^64^. For experiments regarding uninjured and regenerating muscles, homozygous *Rosa26-LSL-H2B-GFP* mice with heterozygous *HSA-Cre* were used. For *Calb1-IRES-Cre* and *Mdx* experiments, the *H2B-GFP* allele was heterozygous. Nuclei were isolated from muscle of 2.5 months old mice. Genotyping was performed as instructed by the Jackson laboratory. For genotyping of the *Rosa26-Lsl-H2B-GFP* reporter, the following primers were used. Rosa4; 5’-TCA ATGGGCGGGGGTCGTT-3’, Rosa10; 5’-CTCTGCTGCCTCCTGGCTTCT-3’, Rosa11; 5’-CGAGGCGGATCACAAGCAATA-3’. Ebf1 mutants^65^ and Dysferlin mis-sense mutants^66^ were described. To induce muscle injury, 30µl of cardiotoxin (10 µM, Latoxan, Porte les Vaence, France) was injected into the tibialis anterior (TA) muscle. Further information on mouse conditions are summarized in Supplementary Table 5.

### Preparation of tissue sections

Freshly isolated TA muscles were embedded in OCT compound and processed as previously described^67^. Frozen tissue blocks were sectioned to 12-16 µm thickness, which were stored at - 80°C until future use.

### Single-molecule FISH (RNAscope)

Otherwise specifically indicated as ‘coventional FISH’ in the figure legends, all the FISH experiments were single molecule FISH using RNAscope. RNAscope_V2 kit was used according to manufacturer’s instructions (ACD/bio-techne). We used Proteinase IV. When combined with antibody staining, after the last washing step of RNA Scope, the slides were blocked with 1% horse serum and 0.25% BSA in PBX followed by primary antibody incubation overnight on 4°C. The subsequent procedures were the same as regular immunohistochemistry. Slides were mounted with Prolonged Antifade mounting solution (Thermofisher). The following probes were used in this study; Smox (559431), Mettl21c (566631), Pdk4 (437161), Ttn (483031), Rian (510531); also synthesized in c2, Vav3 (437431), Col6a3 (552541), Egr1 (423371), Gm10800 (479861), Myh2 (452731-c2) and Calcrl (452281). The following probes were newly designed; Arrdc2 (c1), Cish (c1), Nmrk2 (c1), Slc26a2 (c1), Prkar1a (c1), Tigd4 (c1), Muc13 (c1), Gssos2 (c1), Flnc (c1), Klhl40 (c1), Myh1 (c1), Col22a1 (c2), Ufsp1 (c2), Gm10801 (c2), Xirp1 (c2), Fst1l (c2), human Flnc (c2), Ablim2 (c3), Myh2 (c3) and human Xirp1 (c3).

### Preparation of conventional FISH probes

Probes of 500-700 bp length spanning exon-exon junction parts were designed using software in NCBI website. Forward and reverse primers included Xho1 restriction site and T3 promoter sequence, respectively. cDNA samples prepared from E13.5-E14.5 whole embryos were used to amplify the target probes using GO Taq DNA polymerase (Promega). PCR products were cloned into pGEM-T Easy vector (Promega) according to manufacturer’s guideline, and the identity of the inserts was confirmed by sequencing. 2 µg of cloned plasmid DNA was linearized, 500 ng DNA was subjected to *in vitro* transcription with T3 polymerase and DIG- or FITC-labeled ribonucleotides (All Roche) for 2 hours at 37°C. Synthesized RNA probes were purified using RNeasy kit (Qiagen). Probes were eluted in 50 µl ultrapure water (Sigma), and 50 µl formamide was added. We checked the RNA quality and quantity by loading 5 µl RNA to 2% agarose gel. Until future use, probes were stored in -80°C. The annealing sequences of the FISH probes used in this study are available in Supplementary Table 6.

### Conventional FISH and immunohistochemistry

Basic procedure for conventional FISH was described before^67^ with minor modifications to use fluorescence for final detection. After hybridizing the tissue sections with DIG-labeled probes, washing, RNase digestion and anti-DIG antibody incubation, amplification reaction was carried out using TSA-Rhodamine (1:75 and 0.001% H2O2). After washing, slides were mounted with Immu-Mount (Thermo Scientific). When applicable, GFP antibody was added together with anti-DIG antibody.

When conducting double FISH, the tissue was hybridized with DIG- and FITC-labeled probes, after detection of the DIG signal, slides were treated with 3% H2O2 for 15 minutes and then with 4% PFA for one hour at room temperature to eliminate residual peroxidase activity. The second amplification reaction was performed using anti-FITC antibody and TSA-biotin (1:50), which was visualized using Cy5-conjugated anti-streptavidin.

Antibodies used for this study were: GFP (Aves labs, 1:500), Col3 (Novus, 1:500), ColIV (Millipore, 1:500), CD31-PE (Biolegend, 1:200), F4/80 (Abcam ab6640, 1:500) and Laminin (Sigma L9393, 1:500). For Pax7, we used an antigen retrieval step. For this, after fixation and PBS washing, slides were incubated in antigen retrieval buffer (diluted 1:100 in water; Vector) pre-heated to 80°C for 15 minutes. Slides were washed in PBS and continued at permeabilization step. Cy2-, Cy3- and Cy5-conjugated secondary antibodies were purchased from Dianova.

### FISH experiments using isolated single fibers

We isolated single extensor digitorum longus (EDL) muscle fibers as described before^68^. Isolated EDL fibers were immediately fixed with 4% PFA and were subjected hybridization in 1.5 ml tubes. After DAPI staining, fibers were transferred on slide glasses and mounted. Spindle fibers were vulnerable to collagenase treatment. Thus, we pre-fixed the EDL tissue and peeled off spindles under the fluorescent dissecting microscope (Leica).

### Acquisition of fluorescence images

Fluorescence was visualized by laser-scanning microscopy (LSM700, Carl-Zeiss) using Zen 2009 software. Images were processed using ImageJ and Adobe Photoshop, and assembled using Adobe Illustrator.

### Cell culture

C2C12 cell line was purchased from ATCC, and cultured in high glucose DMEM (Gibco) supplemented with 10% FBS (Sigma) and Penicilllin-Streptomycin (Sigma). To engineer C2C12 cells with doxycycline (Sigma) inducible Ebf1, mouse Ebf1 cDNA (Addgene) was cloned into pLVX Tet-One Puro plasmid (Clontech), packaged in 293T cells (from ATCC) using psPAX2 and VsvG (Addgene), followed by viral transduction to C2C12 cells with 5 µg/µl polybrene (Millipore). Transduced cells were selected using 3 µg/µl puromycin (Sigma).

### Western blotting

Cell pellets were resuspended in NP-40 lysis buffer (1% NP-40, 150 mM NaCl, 50 mM Tris-Cl pH 7.5, 1 mM MgCl2 supplemented with protease (Roche) and phosphatase (Sigma) inhibitors), and incubated on ice for 20 minutes. Lysates were cleared by centrifuging in 16,000 g for 20 min at 4°C. Protein concentration was measured by Bradford assay (Biorad), and lysates were boiled in Laemilli buffer with beta-mercaptoethanol for 10 minutes. Denatured lysates were fractionated by SDS-PAGE, transferred into nitrocellulose membrane, blocked with 5% milk and 0.1% Tween-20 in PBS, and incubated overnight in 4°C with primary antibodies diluted in 5% BSA and 0.1% Tween-20 in PBS. After three times washing with PBST, membranes were incubated with secondary antibodies diluted in blocking solution for one hour at room temperature. After PBST washing, membranes were developed with prime ECL (Amarsham). The antibodies used for this study were β-actin (Cell Signaling, 1:1000), ColVI (Abcam, 1:2000) and Ebf1 (1:1000).

### RT-qPCR

Cell pellets were resuspended in 1 ml Trizol (Thermofisher). RNA was isolated according to manufacturer’s guideline. 1 µg of isolated RNA and random hexamer primer (Thermofisher) were used for reverse transcription using ProtoSciprt II RT (NEB). Synthesized cDNA was diluted five times in water, and 1 µl was used per one qPCR reaction. qPCR was performed using 2X Syber green mix (Thermofisher) and CFX96 machine (Biorad). We used β-actin for normalization. Primers were selected from the ‘Primer bank’ website. The RT-qPCR primers used in this study are available in Supplementary Table 6.

### Human biopsies

Human muscle biopsy specimens were obtained from M. vastus lateralis. We selected wheelchair-bound patients with confirmed *DYSTROPHIN* or *DYSF* mutations and severe dystrophic myopathological alterations defined by histology of biopsies. The exact mutations are indicated in the corresponding Figure legends. The tissues were snap frozen under cryoprotection. Research use of the material was approved by the regulatory agencies (EA1/203/08, EA2/051/10, EA2/175/17, Charité Universitätsmedizin Berlin, Germany). Informed consent was obtained from the donors.

### Reporting summary

Further information on research design is available in the Nature Research Reporting summary linked to this article.

## Data availability

The next-generation sequencing datasets generated in this study are available in the ArrayExpress under accession numbers E-MTAB-8623. The raw DGE count matrix in loom format is available in http://bimsbstatic.mdc-berlin.de/akalin/MyoExplorer/mm10_UMI.loom. Preliminary DGE matrix as a Seurat object is available in http://bimsbstatic.mdc-berlin.de/akalin/MyoExplorer/mm10.Seurat.RDS. Also, we provide an interactive webpage where the users can explore their gene of interest (https://shiny.mdc-berlin.net/Myocyte_scNuc/). Uncropped Western blot images are provided in Supplementary Figure 14. The data that support the findings of this study are available from the corresponding authors upon reasonable requests.

## Code availability

The codes used in this study are accessible in github server (https://github.com/BIMSBbioinfo/MyoExplorer).

## Acknowledgements

We thank Dr. K. Song for advice and protocols on snRNAseq and Dr. T. Muller for critical reading of the manuscript, C. Paeseler and P. Stallerow for help with animal care (all MDC), as well as the MDC core facilities for flow cytometry (led by Dr. Hans-Peter Rahn) and next generation sequencing (led by Dr. Sascha Sauer). We are grateful to Dr. Martyn Goulding (Salk institute) for providing the H2B-GFP reporter mouse line, and Drs. M. Derecka and R. Grosschedl (MPI for Immunobiology and Epigenetics) for providing Ebf1 tissues and reagents. This work was supported by the Helmholtz association (C.B) and AVH postdoctoral fellowship (M.K).

## Author contributions

M.K and C.B conceived the work and designed the project. M.K. led the project, performed the experiments and analyzed the data. V.F and A.A performed the bioinformatics analysis. B.B contributed to the experiments, especially generated the sequencing library. E.D.L also helped with library generation. V.S and S.S collected and prepared human patient biopsies and *dysferlin* mutant mouse samples. M.K and C.B wrote the manuscript with comments by all authors.

