## Supplementary figures for "Single-nucleus transcriptomics reveals functional compartmentalization in syncytial skeletal muscle cells"

**Supplementary Figures Legends.**

**Supplementary Figure 1. Validation of myonuclear labeling by H2B-GFP.** **a**, Efficient labeling of myonclei by H2B-GFP in uninjured and regenerating muscles 7 days after cardiotoxin (CTX) injury (7 d.p.i.). Note the centralized nuclei in 7 d.p.i. Scale bar, 30 μm. **b**, Efficient labeling of myonuclei in isolated single EDL fiber. DAPI was used as a counterstain to identify all nuclei. Scale bar, 100 μm. **c**, Specificity of H2B-GFP expression in myofibers. Double immunoflourescence of GFP and markers of endothelia, Schwann cells, macrophages and muscle stem cells demonstrates the absence of H2B-GFP in these cell types. Scale bar, 50 μm. **d**, Sorting strategy of myonuclei isolation. After gating based on FSC/SSC (P1), doublets were removed (P2 and P3), DAPI positive nuclei were gated (P4), and DAPI/GFP double positive nuclei were sorted (P5). Percentages of gated nuclei (relative to total events) at each step are indicated. DAPI/GFP images before and after sorting are shown on the right. Scale bar, 30 μm.

**Supplementary Figure 2. Quality control plots for the datasets used in this study.** **a**, Number of detected UMIs per nucleus when considering both exonic and intronic reads (left) or only exonic reads (right). **b,** Number of detected genes per nucleus when considering genes detected in at least 5 nuclei. **c,** Mitochondrial reads percentage thresholds for each batch. For all box plots, the center lines indicate median, the end points of the boxes indicate low and high quartiles, and the whiskers indicate minimum and maximum values. Also see Supplementary Table 5 for additional quality control information.

**Supplementary Figure 3. *Myh2* can be co-expressed with *Myh1*, but not with *Myh4*.** Single molecule FISH of *Myh1, Myh2* and *Myh4* in uninjured TA muscle. The probe against *Myh2* detected pre-mRNA and therefore appears as puncta; the probes against Myh1 and Myh4 detected processed mRNA located mainly in the cytoplasm. (Most right) Overlay of Myh2 and Myh4 does not show co-expression of these two Myosins. (Middle) Myh1 and Myh2 can be co-expressed in the same fiber, which are indicated in arrows. Scale bar, 30 μm. Expression patterns were validated in more than 3 individuals.

**Supplementary Figure 4. Genes with dynamic expression profile during regeneration. a**, Violin plots showing the expression level of indicated genes in uninjured, 7 d.p.i. and 14 d.p.i. muscles. **b,** RT-qPCR analysis of indicated genes using isolated GFP+ nuclei at different time points. Error bars indicate S.E.M. Two-tailed unpaired student's t-test (n=3 for each time point). *, $p < 0.05$. **, $p < 0.01$. ***, $p < 0.001$. **c,** Single molecule FISH of indicated genes at different time points. Scale bar, 50 μm. Expression patterns were validated in 2 or more individuals.

**Supplementary Figure 5. Expression of *Tigd4*, a marker of MTJ-A, in uninjured TA muscle.** Tile scan image of single molecule FISH against *Tigd4*. DAPI was used as a counterstain to identify all nuclei. The insets indicate the MTJ area that is magnified in the right panels. Expression patterns were validated in more than 3 individuals. Scale bars, 200 μm (for the entire tile scan) and 100 μm (for the insets).

**Supplementary Figure 6. FISH validation of the expression of marker genes identified in the MTJ-B cluster.** Conventional FISH (*Col6a3* or *Pdgfrb*) and GFP immunofluorescence at the MTJ in uninjured (top two rows) or 14 d.p.i. TA muscle expressing H2B-GFP in myonuclei (middle two rows). T indicates tendon and M myofiber; arrowheads point to myonuclei that co-express marker genes and GFP. The bottom two rows show regions distal to the MTJ in uninjured muscle where expression of marker genes is also observed in GFP-negative nuclei. Note that due to high temperatures used during the conventional FISH (but not single molecule FISH) procedure, GFP signals are dampened and not observed in all myonuclei. Expression patterns were validated in 2 or more individuals. Scale bar, 50 µm.

**Supplementary Figure 7. Quantification of repeat elements in the snRNAseq dataset.** Unsupervised hierarchical clustering of the repeat element expression profile across nuclear populations. The most frequently expressed 16 repeat elements are presented. Note the strong enrichment of srpRNA in Gssos2+ nuclei.

**Supplementary Figure 8. *Rian* and *Gssos2* are expressed in distinct nuclear populations. a,** Single molecule FISH of *Rian* and *Gssos2* in uninjured TA muscle shows that *Rian* expressing nuclei (asterisk) and *Gssos2* expressing nuclei (arrows) are distinct. **b,** Single molecule FISH of *Rian*/*Ufsp1* and *Gssos2*/*Ufsp1* shows that *Rian* and *Gssos2* expressing nuclei (arrows) are distinct from Ufsp1 expressing nuclei. Expression patterns were validated in 2 or more individuals. Scale bars, 30 µm.

**Supplementary Figure 9. Re-clustering of the bulk myonuclei identified in Figure 1.** Labeling of different time points (left) and expression profile of *Ttn* (right).

**Supplementary Figure 10. Further characterization of nuclear subtypes in *Mdx* muscle. a,** Expression of *Ttn* and *Dystrophin*. Note that *Dystrophin* expression is homogenous across all nuclear populations. **b,** MA plot of enriched genes in the 'dying fiber' cluster in Figure 4a; Exp., expression. **c**, Co-expression of indicated ncRNAs in a subset of myonuclei. Scale bar, 30 µm. **d**, Fibers containing such nuclei are infiltrated with H2B-GFP negative cells in c. Scale bar, 30 µm. **e,** Location of MTJ-A in the *Mdx* UMAP map. Scales indicate gene signature scores of MTJ-A nuclei. **f,** Single molecule FISH of NMJ markers (*Ufsp1* and *Prkar1a*) or MTJ-A markers (*Tigd4* and *Col22a1*) in control and *Mdx* muscle. Note that the strict co-expression of *Prkar1a* and *Ufsp1* is weakened in *Mdx* muscle. In contrast, *Tigd4* and *Col22a1* are strongly co-expressed in *Mdx* muscle. Scale bar, 50 µm.

**Supplementary Figure 11. Myonuclei expressing a repair gene signature in Mdx and Dysferlinopathy muscles. a,** Single molecule FISH images for *Flnc*/*Xirp1* in wild-type mouse or muscle of a healthy human donor. These FISH experiments were conducted at the same time as those shown in Figures 4f and 4g. **b**, Co-expression of indicated marker genes in muscle of *Mdx* mice. **c**, Co-expression of *Flnc* and *Xirp1* in the muscle of mice carrying a mis-sense mutation (L1360P) in the *Dysferlin* gene. **d**, Co-expression of *FLNC* and *XIRP1* in a muscle biopsy of a human patient carrying a homozygous mutation in the *DYSFERLIN* gene (*DYSF* c.4872delG);

nuclei co-expressing these genes were also identified in a second patient. Scale bars, 50 µm. Expression patterns in mouse tissues were validated in 2 or more individuals.

**Supplementary Figure 12. Validation of nuclear compartments in the muscle spindle. a,** Two different protocols were used to isolate spindle myonuclei, and this does not bias their distribution in the UMAP plot. See methods for further details of these protocols. **b,** Expression of indicated marker genes by single molecule FISH in cross-sections of TA muscle expressing H2B-GFP after *Calb1-Ires-Cre* recombination. Scale bars, 10 µm. Expression patterns were validated in 2 or more individuals.

**Supplementary Figure 13. Loss of Ebf1 does not affect expression of marker genes in NMJ, MTJ-A and Rian+ nuclei.** Single molecule FISH of indicated marker genes in TA muscle of control or *Ebf1* mutant mice. Expression patterns were validated in 3 individuals. Scale bar, 30 µm.

**Supplementary Figure 14. Uncropped Western blot images for figure 6c.**

### Supplementary Fig. 1

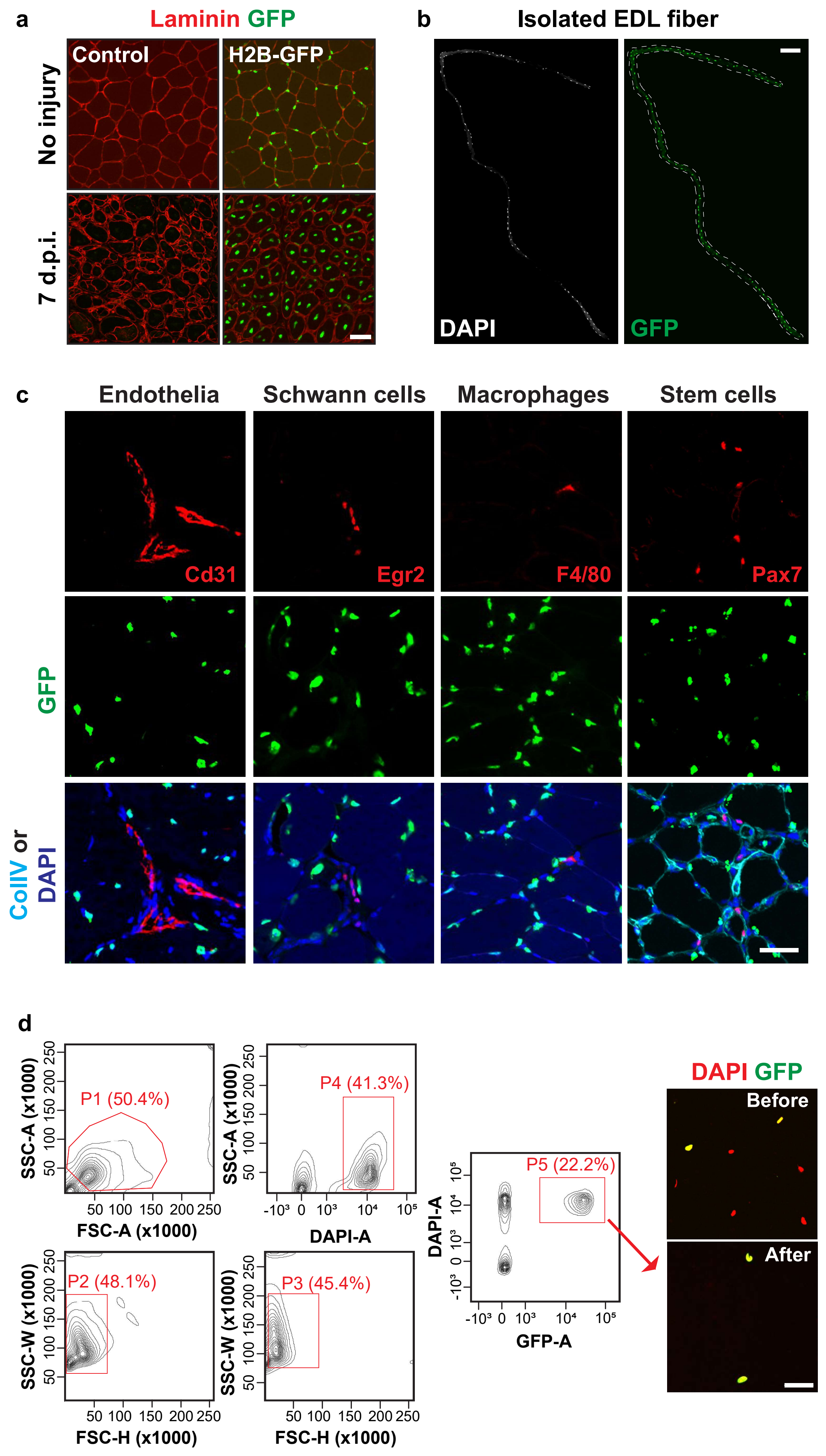

### Supplementary Fig. 2

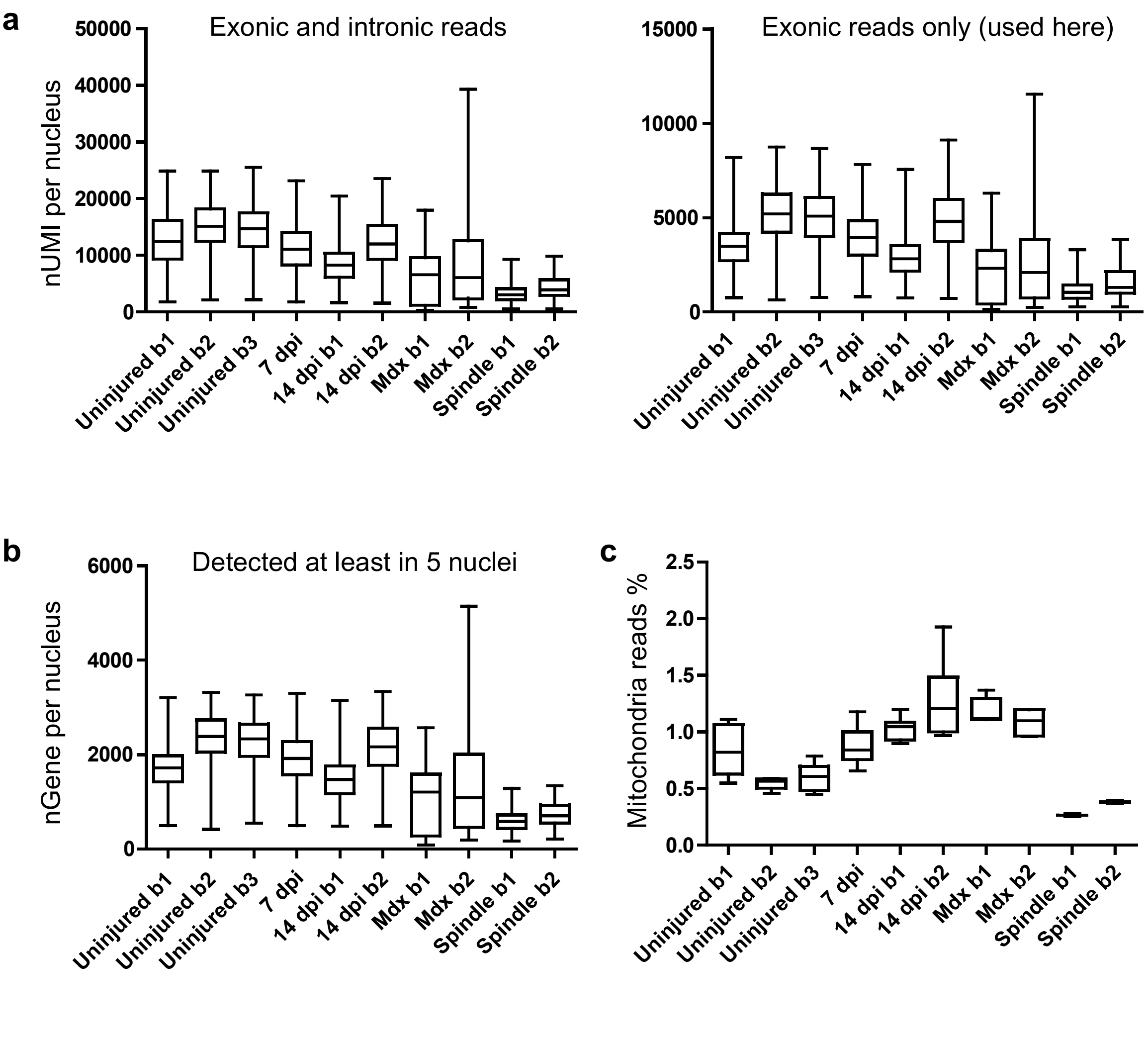

**Supplementary Fig. 3**

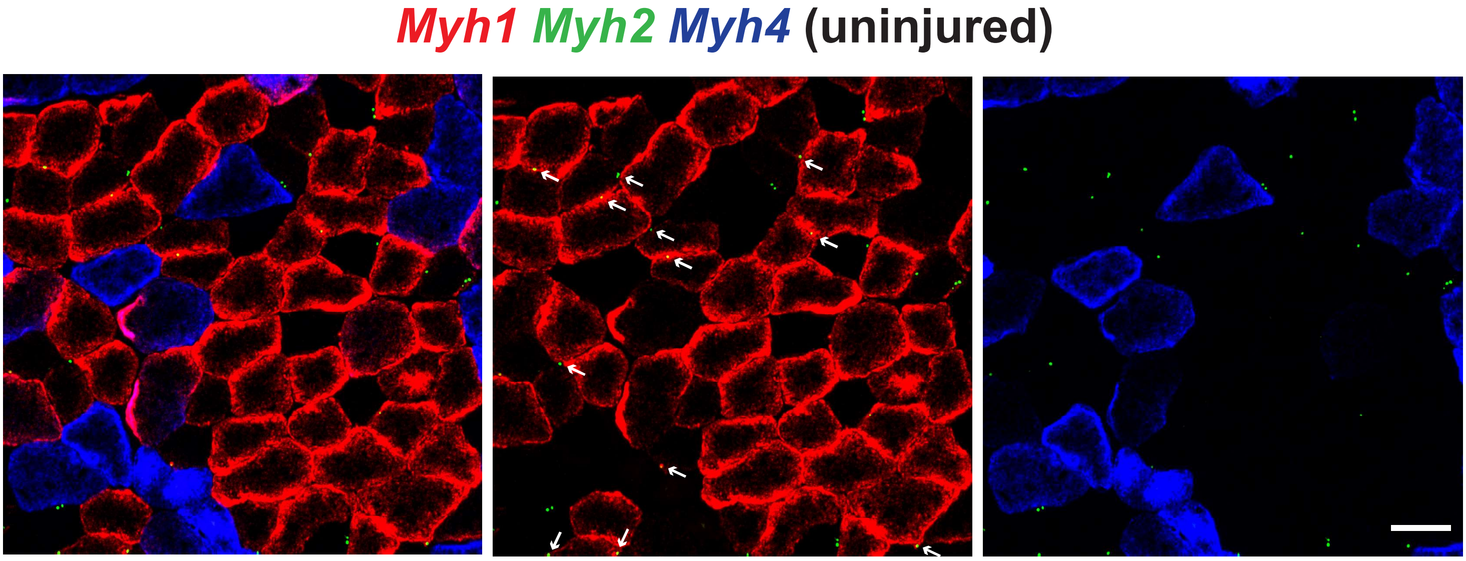

### Supplementary Fig. 4

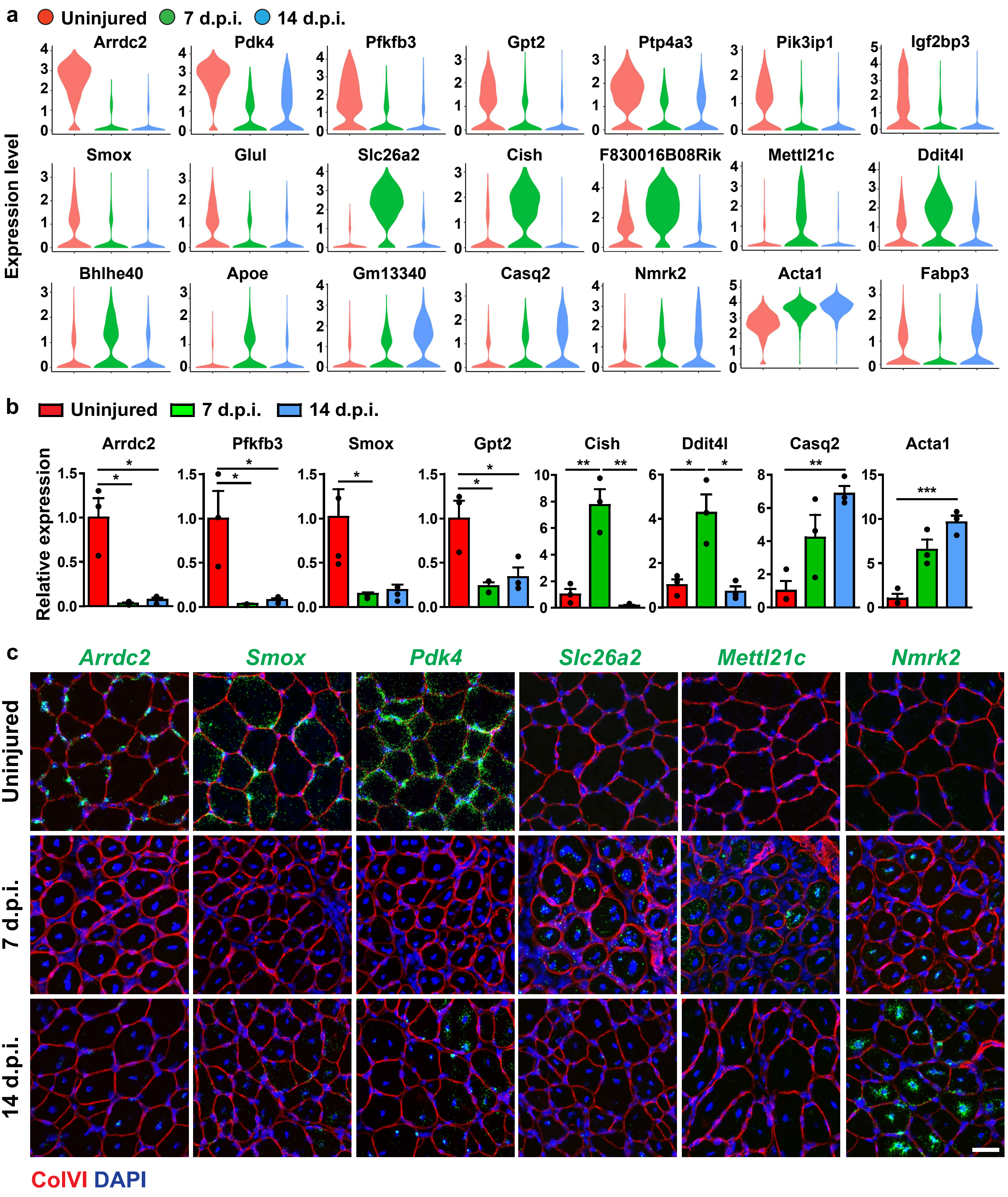

**Supplementary Fig. 5**

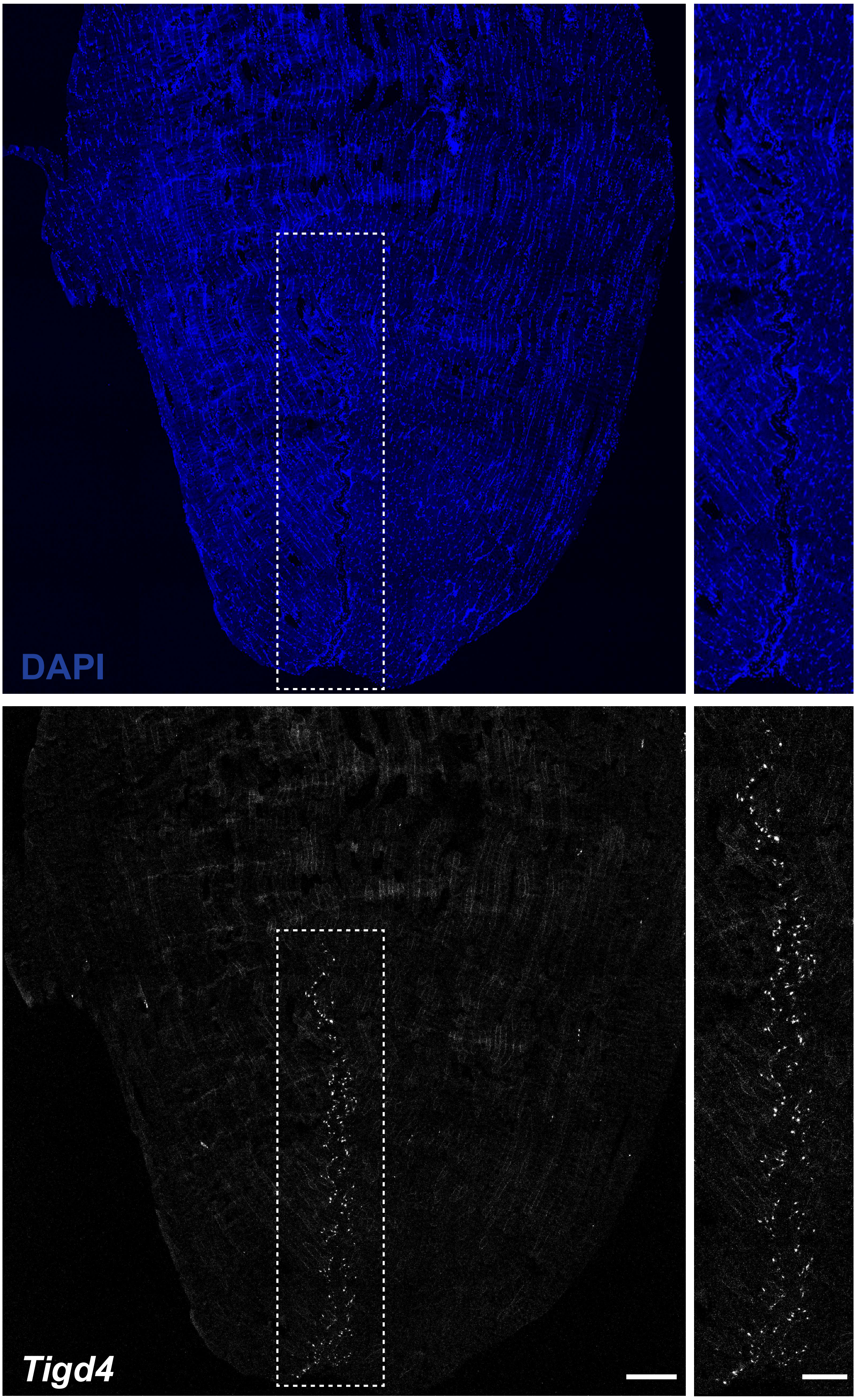

### Supplementary Fig. 6

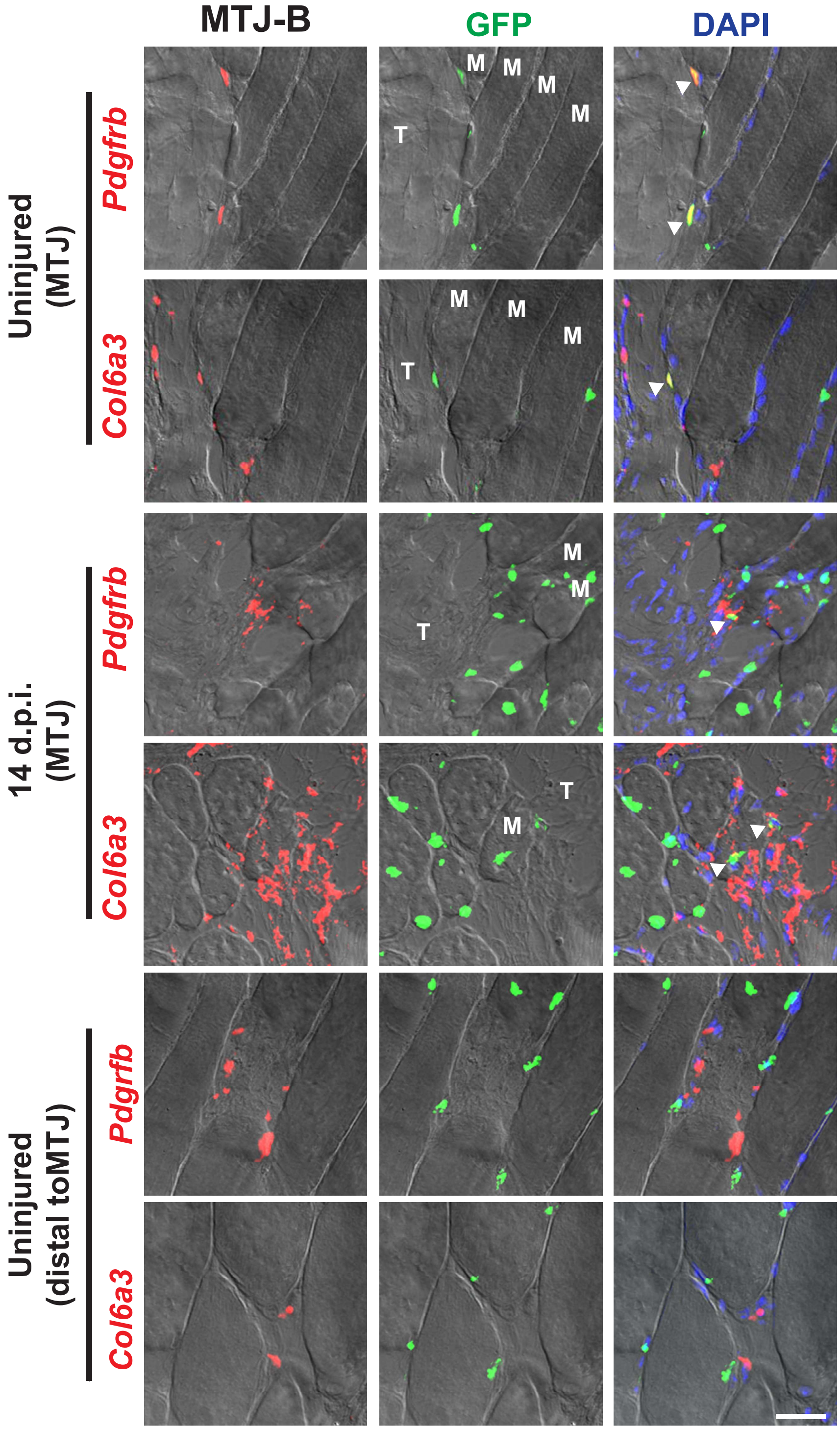

**Supplementary Fig. 7**

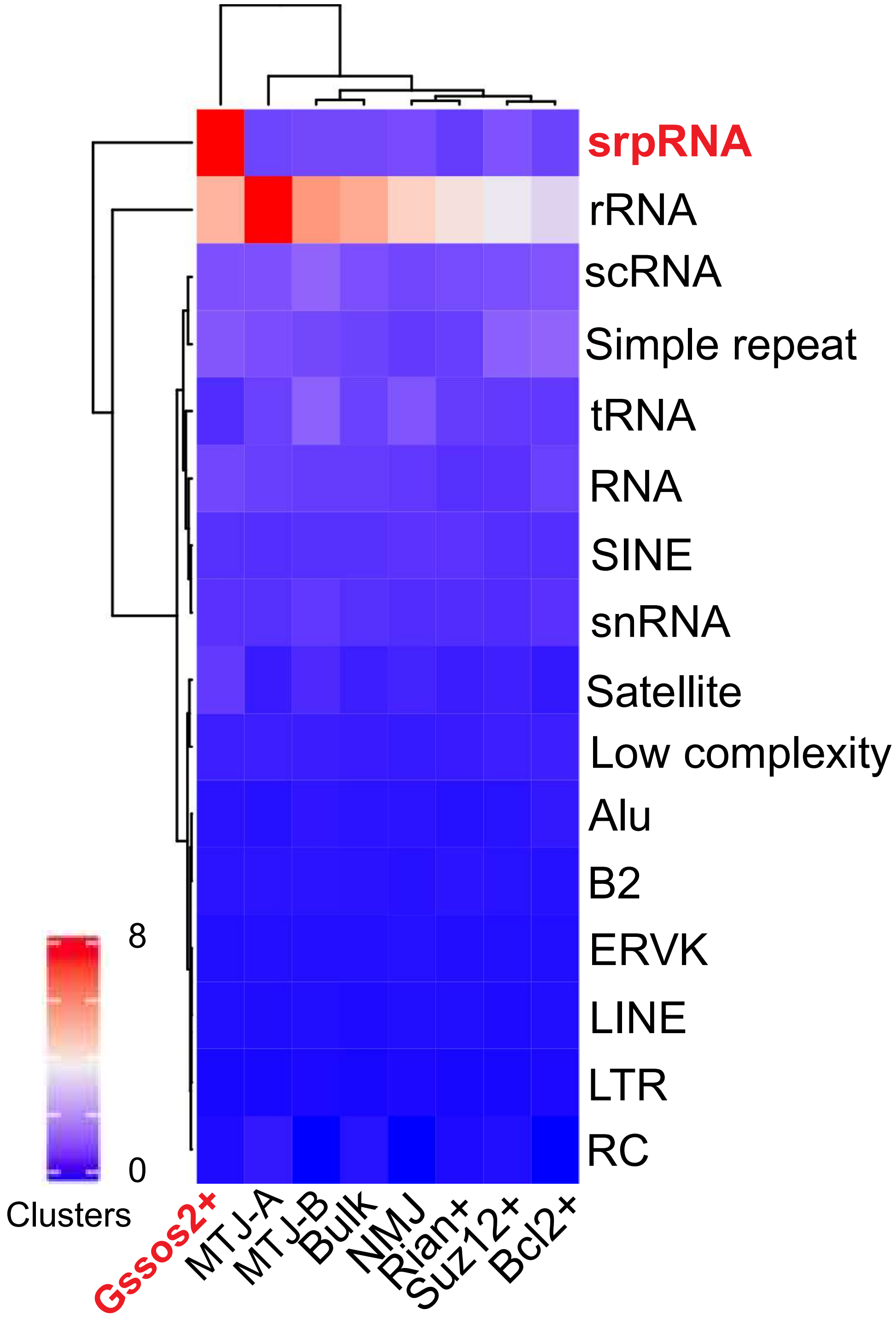

**Supplementary Fig. 8**

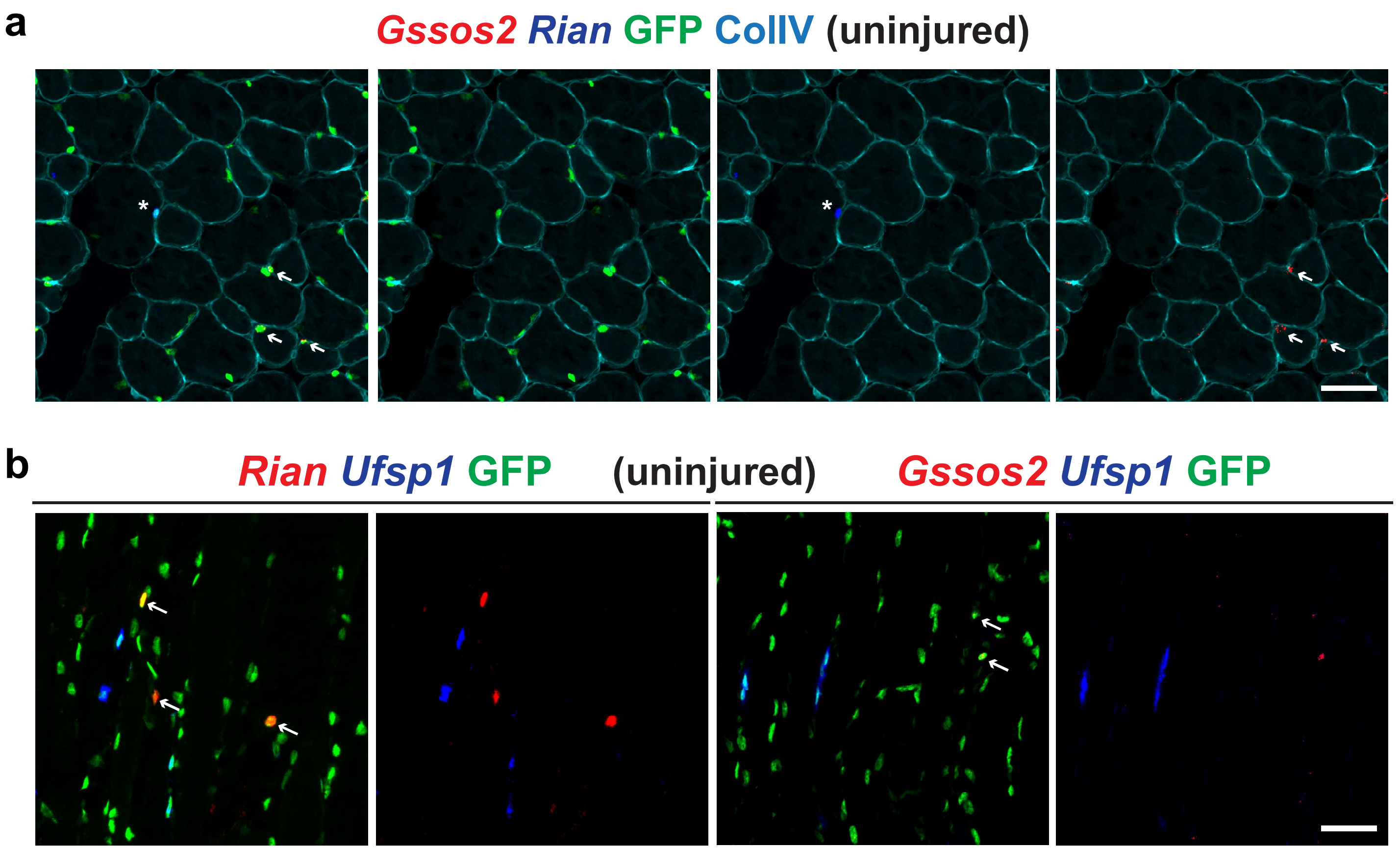

**Supplementary Fig. 9**

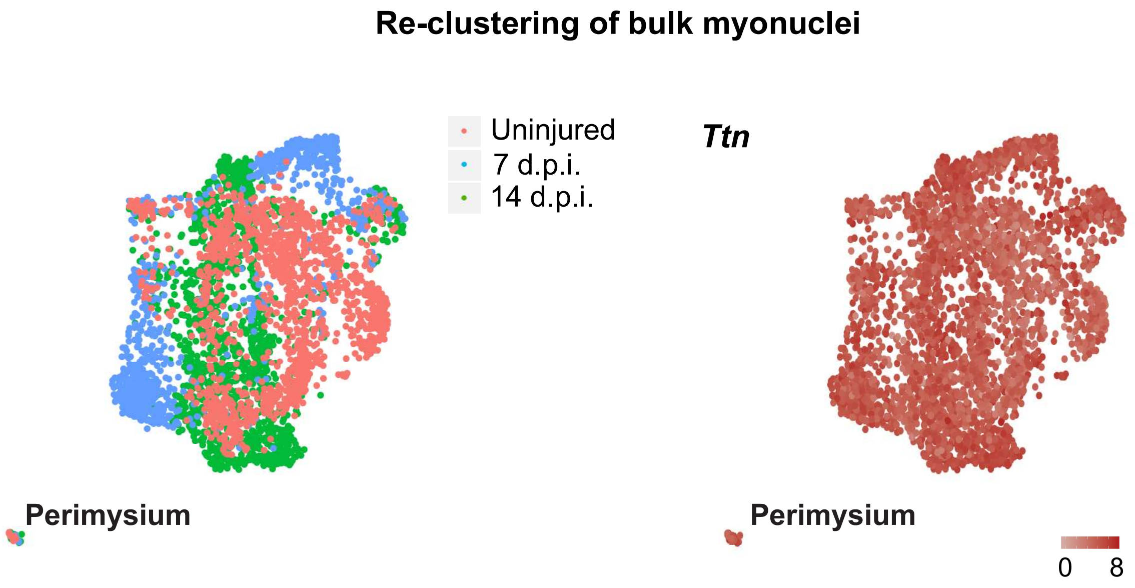

**Supplementary Fig. 10**

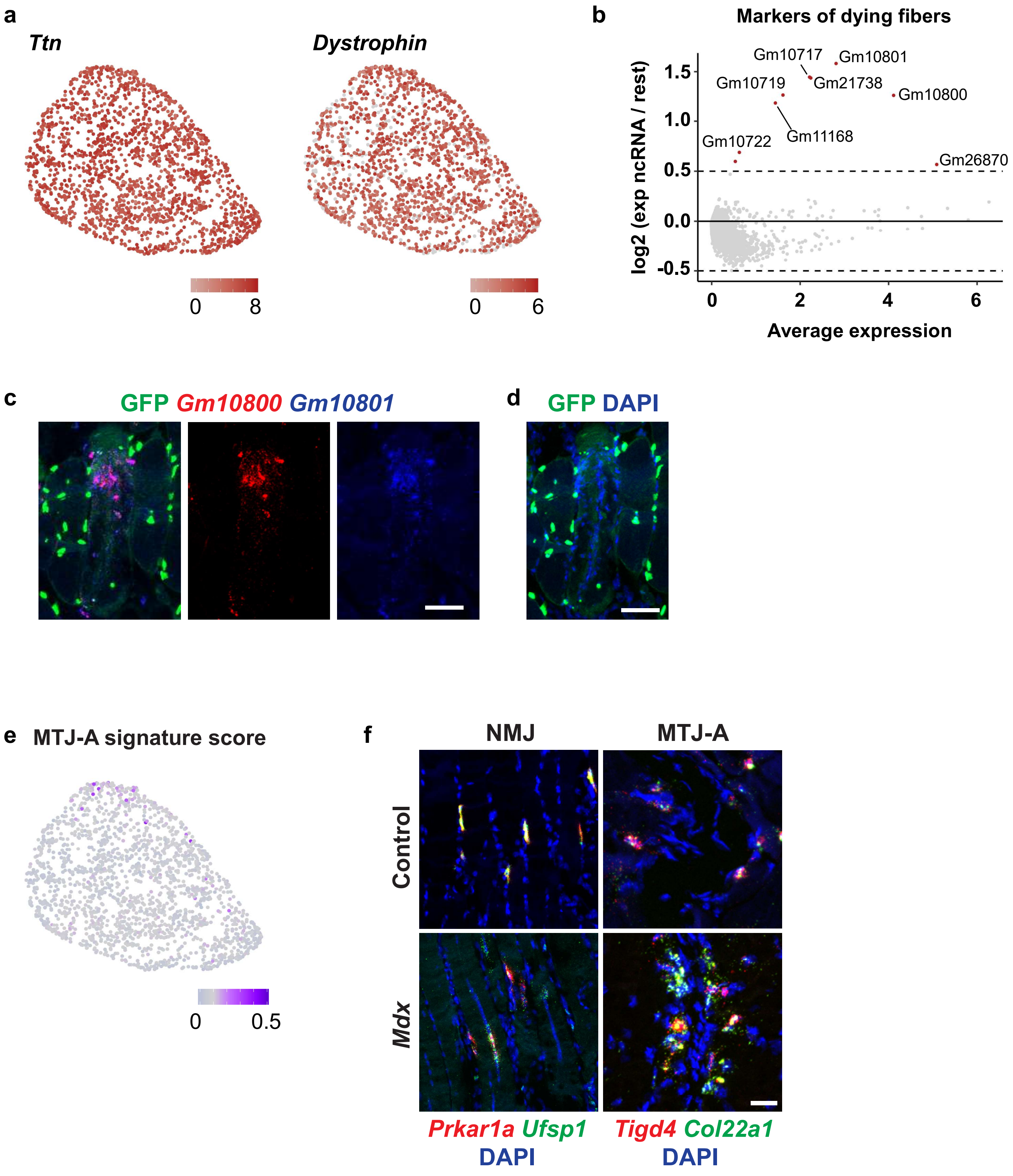

### Supplementary Fig. 11

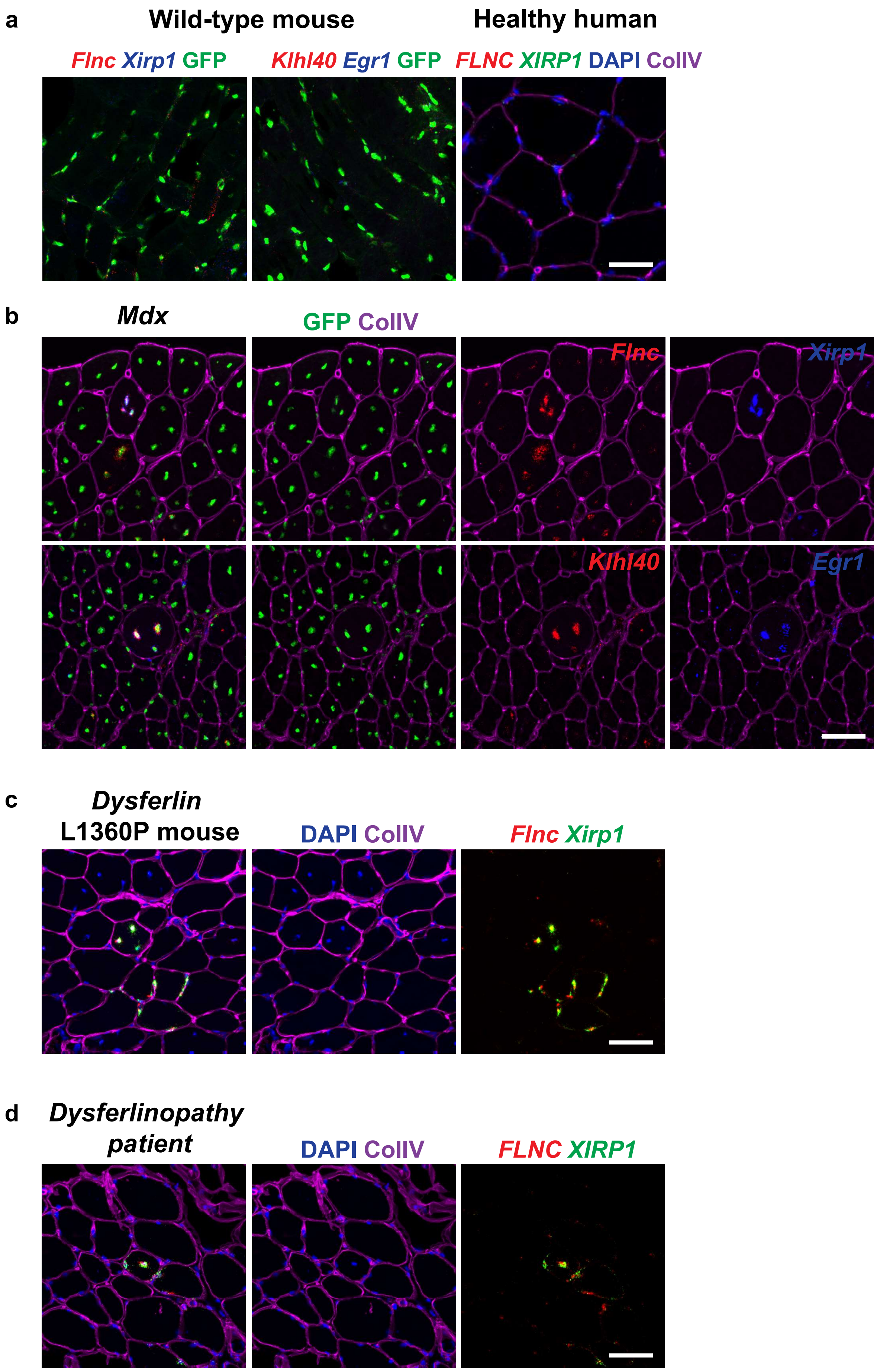

### Supplementary Fig. 12

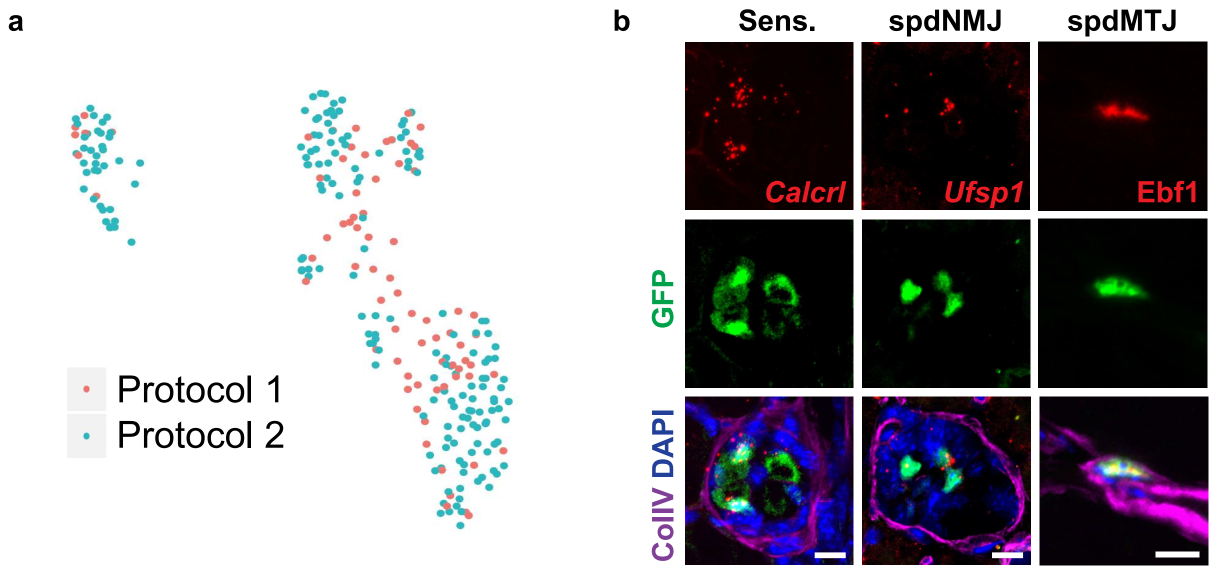

**Supplementary Fig. 13**

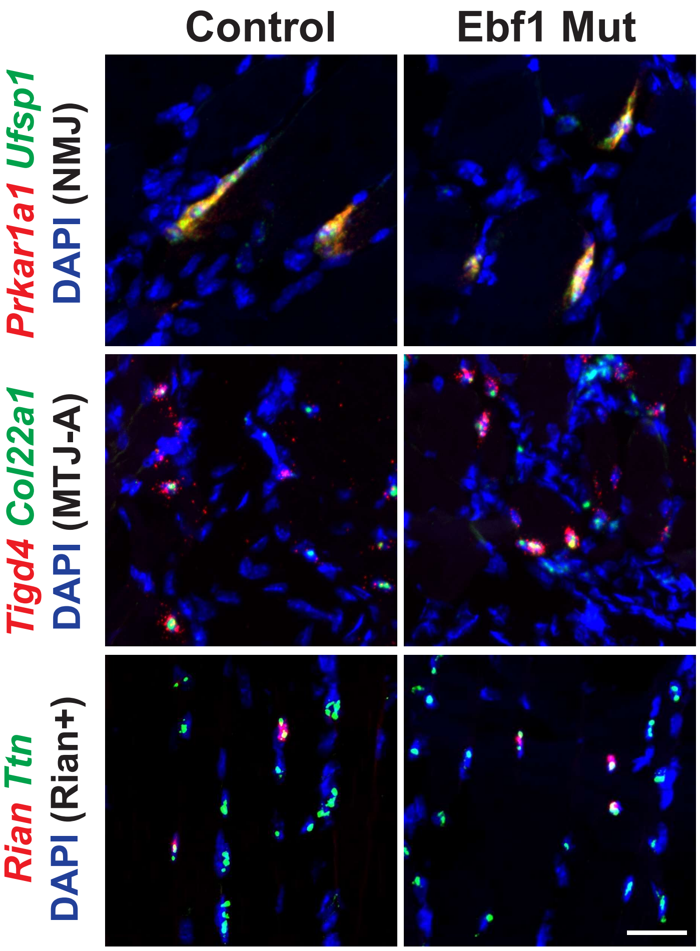

**Supplementary Fig. 14**

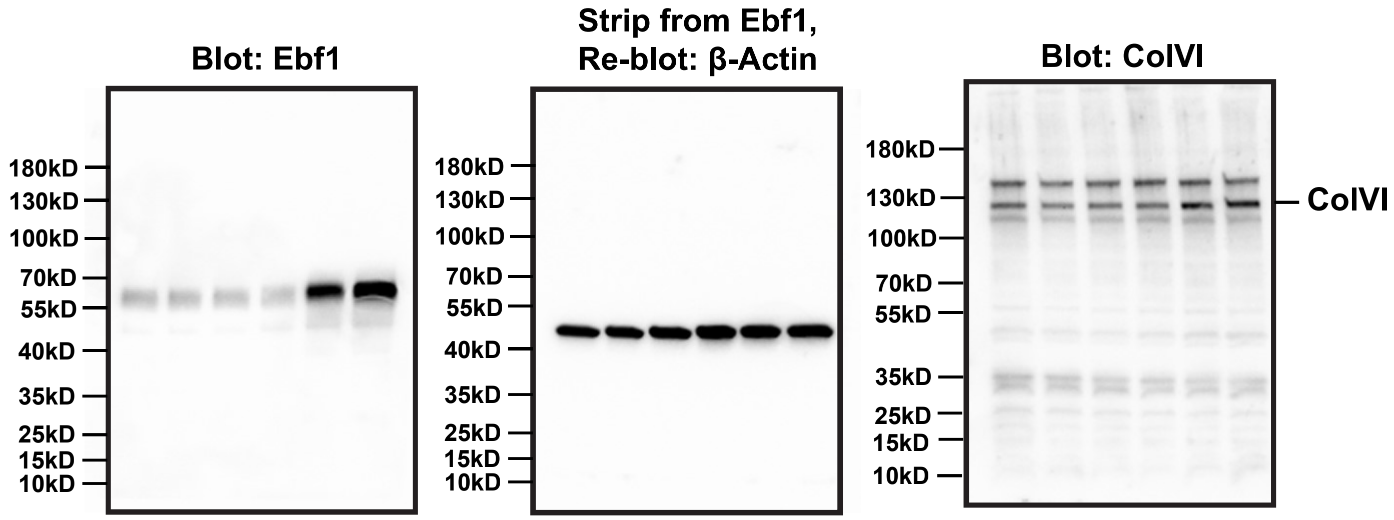
